# Sphingolipids integrate TORC2 with TORC1 through vacuolar liquid-ordered domain formation

**DOI:** 10.64898/2026.06.03.729730

**Authors:** Hiroto Denda, Katsuki Eto, Kazue Nishiyama, Ritsuko Hagiwara, Tatsuki Okano, Shunsuke Yoshizumi, Ayaka Hirota, Kouta Nakazono, Yukari Yabuki, Akira Okano, Yuichiro Sekikawa, Toshi Shimamoto, Kazuki Hanaoka, Keiko Sakuragi, Philipp Schlarmann, Atsuko Ikeda, Koji Masumura, Masaki Mizunuma, Naohiro Ise, Muneyoshi Kanai, Haruyuki Iefuji, Aswin T. Srivatsav, Pranav Adhyapak, Lydia Mathew, Shobhna Kapoor, Kouichi Funato

## Abstract

The coordination of multiple signaling pathways across different compartments of the endomembrane system is essential for cellular adaptation to the environment; however, the underlying mechanisms remain poorly understood. Previous studies have described the existence of sterol-enriched liquid-ordered (Lo) domains in the vacuole membrane of yeast, which act as platforms for signal initiation. Here, we demonstrate that vacuolar Lo domains serve as signaling platforms for the evolutionarily conserved Target of Rapamycin Complex 1 (TORC1) kinase complex. Notably, we show that TORC2, which is functionally distinct from TORC1 and localized to the plasma membrane, regulates TORC1 activity through sphingolipid metabolism and subsequent vacuolar Lo domain formation. Collectively, our findings reveal a lipid domain-mediated network that integrates two signaling pathways originating from spatially different cellular sites.

**HIGHLIGHTS:**

1. Decreased sphingolipid levels cause characteristic phenotypes that reflect low TORC1 activity.
2. Sphingolipid-enriched vacuolar Lo domains are required for the EGO complex-mediated activation of TORC1.
3. Sphingolipids integrate TORC2 and TORC1 through vacuolar Lo domain formation.
4. Vacuole fusion and acidification control TORC1 activity via sphingolipid metabolism.

## INTRODUCTION

Cells have developed sophisticated and evolutionarily conserved mechanisms to sense and adapt to environmental changes in their surroundings. Because the perception of and responses to environmental changes occur at multiple spatially distinct sites within the cell, cross-organellar communication is key to orchestrating a coherent response. Although many communication pathways, such as kinase signaling, vesicular trafficking, and the formation of membrane contact sites (MCS), have been described in great detail,^1–7^ how these pathways are effectively coordinated with each other remains poorly understood.

Similar to plasma membrane microdomains, often referred to as lipid rafts,^8–11^ microdomains within organelle membranes also provide unique environments that may serve as platforms for signal transduction and membrane sorting. The existence of such micrometer-scale domains within cells remains controversial,^8^ but stable microdomains formed during nutrient starvation or glucose depletion have been reported in yeast vacuoles.^12,13^ Vacuolar microdomains are enriched in sterols, which are phase-segregated from regions containing marker proteins for liquid-disordered (Ld) domains, such as Vph1 and Pho8. In addition, the sterol-rich liquid-ordered (Lo) domains are enriched with sphingolipids, and sphingolipids are essential for vacuole phase separation into Ld and Lo domains.^14^ Previous studies showed that proteins involved in vesicular trafficking, such as Sec18, or Vps4 as well as MCS tethering proteins support the formation of vacuolar microdomains.^12,15^ While the importance of vacuolar Lo domains as platforms for incorporating lipid droplets into the vacuolar membrane during microlipophagy is well known,^13,16^ the roles of Lo domains in signal transduction originating from the vacuolar membrane have not been defined. Prior studies have implicated Tco89, Gtr1, Gtr2, and Ego1 as vacuolar Lo domain-associated proteins.^10,13,17^ Tco89 is a subunit of the Target of Rapamycin Complex 1 (TORC1), one of two TOR complexes (TORC1 and TORC2) with distinct constituent factors, roles, and cellular locations. Gtr1, Gtr2, and Ego1 are subunits of the TORC1 upstream positive regulator exit from the G0 (EGO) complex, which localizes to the vacuolar membrane and binds to TORC1.^5,7^ This association of TORC1 and EGO complex subunits with the Lo domain raises the possibility that vacuolar Lo domains may contribute to the regulation of TORC1-mediated signaling. However, their significance in TORC1 signaling emitted from the vacuolar membrane remains unclear.

In this study, we show that inositolphosphorylceramide (IPC), a yeast complex sphingolipid and component of liquid-ordered (Lo) microdomains, plays a crucial role in maintaining vacuolar morphology and regulating TORC1 signaling. Moreover, we found that binding of the EGO complex to the vacuolar Lo domain is required for IPC-mediated TORC1 activation. Our data establish that the vacuolar Lo domain functions as a site that integrates TORC1 and TORC2 signaling.

## RESULTS

### Genomic and cellular biological analyses suggest a significant involvement of sphingolipids in the regulation of vacuole morphology and autophagy

To identify the biological processes governed by sphingolipids in yeast, we performed a chemical genetic screen using the *S. cerevisiae* haploid deletion mutant collection (4986 mutants) for genes that modulate the sensitivity of aureobasidin A, an inhibitor of inositolphophoryl ceramide (IPC) synthase, Aur1. The sensitivity of yeast mutants to AbA was assessed by spotting cell suspensions on minimal synthetic defined (SD) medium containing 0.01 μg/ml AbA at 25 °C. This AbA concentration is lower than the limit (0.05 μg/ml) at which wild-type cells exhibit growth defects^18–20^. Screening allowed the identification of 30 genes required for survival under low AbA treatment (Figure S1), and functional categorization showed that approximately two-thirds of these genes were enriched in processes linked to the vacuole, such as homotypic vacuolar fusion, vacuolar acidification, and transport (Figure 1A). We confirmed that deletion of other genes involved in these processes caused hypersensitivity to AbA (Figure S2A, S2B, and S2C). These findings suggest that sphingolipids play a role in vacuole structure and function. In agreement with this, we observed negative genetic interactions of *vps3*Δ (homotypic vacuolar fusion) (Figure S3A), *vma4*Δ (vacuolar acidification) (Figure S3B), *stp22*Δ, *vps36*Δ, *vps20*Δ, or *snf7*Δ (endosomal sorting complex required for transport (ESCRT)) (Figure S3C), with a temperature-sensitive allele of the catalytic component of the IPC synthase *aur1-1*, which was generated in this study. This mutant exhibited the strongest growth phenotype compared to the other alleles (*aur1-6* and *aur1-18*) (Figure S4A and S4B) and caused reduced IPC synthesis and ceramide accumulation even at the permissive temperature of 25 °C (Figure S4C and S4D).

**Figure 1.**
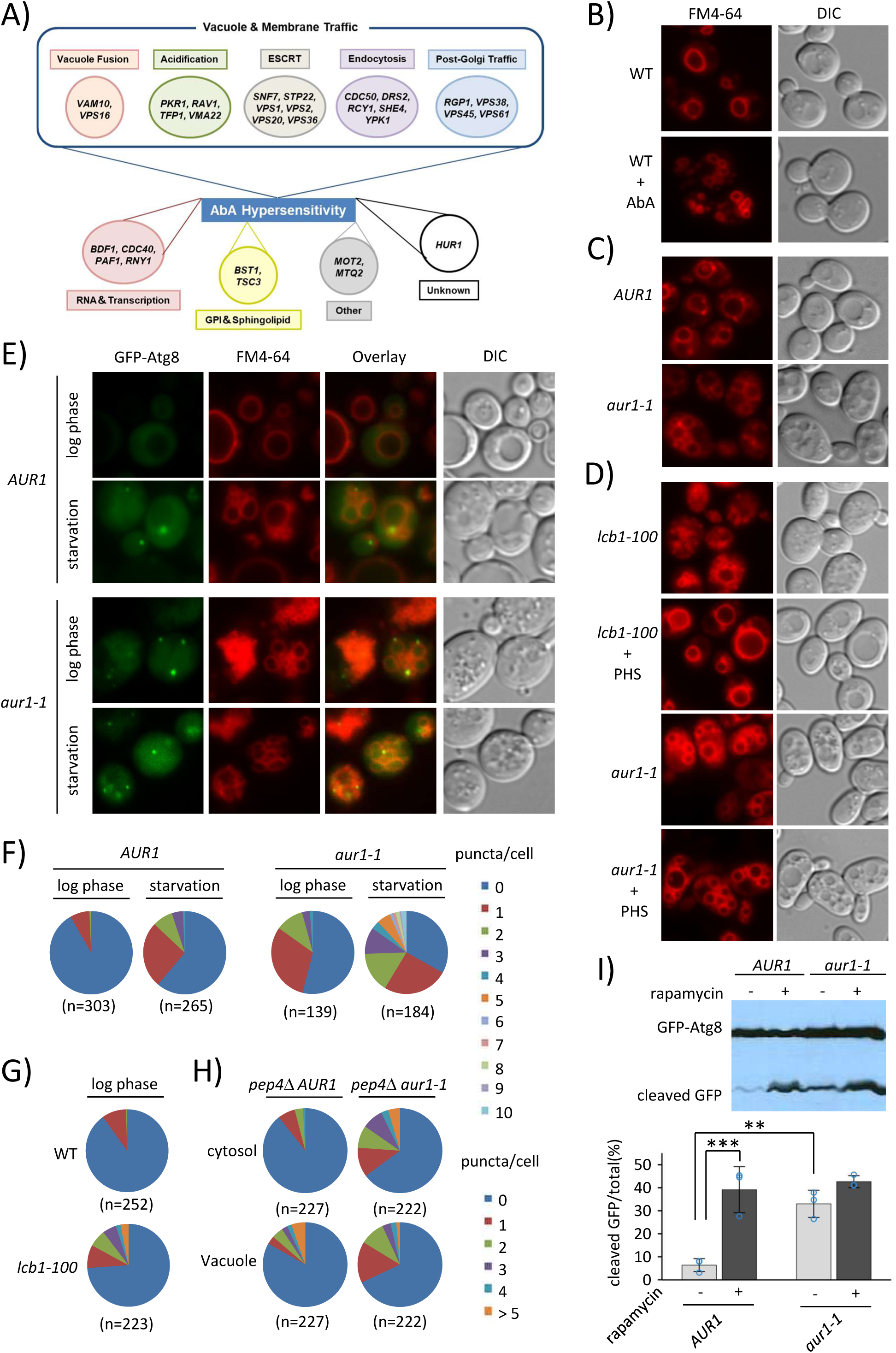
A chemical genomic screen identified functional connections between vacuole function and sphingolipid metabolism, and revealed that reduced sphingolipid levels induce vacuole fragmentation and autophagy. A) Functional classification of genes identified in a chemical genomic screen for genes whose deletion confers sensitivity to aureobasidin A (AbA). B-D) Vacuole morphology of wild-type treated with or without 0.05 μg/ml AbA for 4h (B), *AUR1* and *aur1-1* (C), and *lcb1-100* and *aur1-1* mutants treated with or without 50 μM phytosphingosine (PHS) for 10 h (D). After cells were grown in rich medium (YPD) at 25° °C, the vacuolar membrane was stained with FM4-64 for 2h and visualized using a fluorescence microscope. Fluorescence and DIC images are shown. E) Fluorescence micrographs of GFP-Atg8 in *AUR1* and *aur1-1* mutant cells. The cells expressing GFP-Atg8 were grown exponentially in synthetic dextrose medium (SD), stained with FM4-64 for 1h (log phase), then starved for nitrogen (60 min; starvation), and visualized using a fluorescence microscope. F-H) Quantification of the percentage of cells with GFP-Atg8 dots. The percentage of cells with GFP-Atg8 dots at different numbers shown in (E) was quantified in *AUR1* and *aur1-1* mutant cells (F). At least 100 cells (n=139–303) were counted for each strain. The percentage of cells with GFP-Atg8 dots in wild-type and *lcb1-100* mutant cells (G) and either outside the vacuole (in the cytosol) or in the vacuole of *pep4*Δ *AUR1* and *pep4*Δ *aur1-1* cells (H) grown exponentially is shown. I) Vacuolar degradation of GFP-Atg8 by autophagy. *AUR1* and *aur1-1* cells expressing GFP-Atg8 were grown in YPD and treated with or without 200 nM rapamycin for 60 min, and protein extracts were analyzed by immunoblotting using anti-GFP. Autophagic flux was estimated based on the ratio of cleaved GFP to total GFP (GFP-Atg8 and cleaved GFP) and expressed as a percentage. Average values and standard deviations (SD) were obtained from three independent experiments. **P value for the Tukey-Kramer multiple comparison test < 0.01; ***P < 0.001.

As genetic interactions open up the possibility that sphingolipids play an essential role in the maintenance of vacuole structure and function, we first asked whether AbA treatment and *aur1* mutations impact the number of vacuoles per cell. As shown in Figure 1B, AbA treatment for 4 h increased the number of vacuoles/cell (referred to as vacuole fragmentation) detected using the amphiphilic dye FM4-64, which selectively stains yeast vacuolar membranes. Such fragmented vacuoles were observed in the *aur1-1* mutant at both 25 °C and 37 °C, whereas in the *aur1-6* and *aur1-18* mutants, significant vacuolar fragmentation occurred only at 37 °C (Figure 1C and S5A). Moreover, similar to the *aur1-1* mutant, vacuoles were highly fragmented at 25 °C in the temperature-sensitive *lcb1-100* mutant (Figure 1D), which is defective in serine palmitoyltransferase. This enzyme catalyzes the first step of sphingolipid synthesis, resulting in reduced levels of all sphingolipids, as observed previously.^21^ These results confirm previous reports showing that mutants with lower sphingolipid levels display vacuole morphology defects.^22,23^ Notably, vacuole fragmentation in *lcb1-100* was rescued by exogenous addition of phytosphingosine (PHS), a precursor of phytoceramides that are converted to complex sphingolipids such as IPC, mannosyl IPC (MIPC), and mannosyl-diinositolphosphorylceramide (MIP_2_C).^24,25^ The same was not observed in the *aur1-1* mutant (Figure 1D), suggesting that decreased complex sphingolipid levels are responsible for vacuolar fragmentation. Vacuole fragmentation was not observed in deletion strains lacking genes involved in MIPC (*CSG1* and *CSH1*) and MIP_2_C (*IPT1*) synthesis, as well as sphingolipid hydroxylation (*SUR2* and *SCS7*) (Figure S5B). Therefore, collectively, the data suggest that IPC is crucial for maintaining the number of vacuoles per cell.

Next, we investigated whether a decrease in IPC level affects vacuolar acidification. Vacuolar acidification was assessed by fluorescence staining with quinacrine, which concentrates in acidic lumina.^26,27^ While the *aur1-1* and *lcb1-100* mutants showed typical patterns of quinacrine accumulation in fragmented vacuoles, quinacrine fluorescence was absent in *vma3*Δ cells lacking vacuolar ATPase activity (Figure S6A), suggesting that neither *aur1-1* nor *lcb1-100* is defective in vacuolar acidification. This is consistent with the observation that the *aur1-1* and *lcb1-100* mutants, but not *vma3*Δ, are able to grow at an extracellular pH of 7.5 (Figure S6B), but is inconsistent with a previous study supporting that sphingolipids with a C26 acyl group are required for V-ATPase activity.^28^ Moreover, to test whether sphingolipids are required for ESCRT-dependent protein sorting and endosome-to-vacuole vesicular transport, we analyzed the localization of GFP-Cps1, which is delivered to the vacuolar lumen via the ESCRT pathway^29^ and the kinetics of FM4-64 staining. In both *aur1-1* and *lcb1-100* cells, GFP-Cps1 was properly located in the lumen of the vacuole (Figure S6C), indicating that Cps1 sorting via the ESCRT pathway is normal in mutants defective in sphingolipid synthesis. As a positive control, *vps4*Δcells exhibited mislocalized GFP-Cps1 to the vacuole membrane. The delivery of internalized FM4-64 to the vacuole also remained unaffected in *aur1-1* cells (Figure S6D). Thus, complex sphingolipids are not particularly important for ESCRT-dependent protein sorting and endosome-to-vacuole vesicular transport.

Although ATG genes required for autophagy were not identified in our chemical genetic screen for hypersensitivity to AbA, *AUR1* has been reported to display negative genetic interactions with *ATG1* and *ATG31* and a positive interaction with *ATG15*.^30–32^ Therefore, we investigated whether a functional link exists between sphingolipid metabolism and autophagy. For this, we visualized the localization of GFP-Atg8, a pre-autophagosomal structure (PAS) marker^33^ in mutants defective in sphingolipid synthesis. In *AUR1* cells (*aur1*Δ cells complemented with Sc*AUR1*), GFP-Atg8 puncta were rarely observed in the early logarithmic growth phase, but approximately 40 % of cells displayed GFP-Atg8 puncta upon nitrogen starvation (Figure 1E and 1F). Remarkably, in the *aur1-1* mutant, a larger number of cells with GFP-Atg8 puncta were observed even during the early logarithmic growth phase. A similar observation was made in *lcb1-100* cells (Figure 1G). Moreover, in *pep4*Δ background strains, which are defective in autophagic body degradation, the *aur1-1* mutation increased the appearance of GFP-Atg8 puncta within the vacuole lumen (Figure 1H). Consistent with the imaging data showing that sphingolipid depletion triggers bulk autophagy, the proteolytic generation of free GFP from GFP-Atg8 in the vacuolar lumen was observed in the *aur1-1* mutant (Figure 1I). The production of free GFP in the mutant was similar to that in the control strain treated with rapamycin, which induces autophagy by repressing TORC1 activity. Taken together, these findings suggest that sphingolipids regulate vacuole morphology and autophagy.

### Sphingolipid-deficient cells display phenotypes induced by reduced TORC1 activity

As autophagy is negatively regulated by TORC1 via direct phosphorylation of the autophagy-related protein Atg13,^34^ we investigated whether impaired sphingolipid synthesis affects TORC1 signaling. We analyzed the transcriptomic profiles of AbA-treated and untreated wild-type cells to determine whether AbA induced gene expression changes characteristic of TORC1 inactivation. These included reduced expression of ribosomal protein and RiBi regulon genes and increased expression of *ATG*, *RTG*, *HAP*, and *MSN* targets, as well as genes involved in nitrogen discrimination and glycogen accumulation.^35–42^ As shown in Figure 2A and 2B, the expression of many ribosomal and RiBi proteins was downregulated upon AbA treatment, whereas several genes that are negatively regulated by TORC1 were upregulated. Using *aur1* mutant strains, we confirmed that sphingolipid depletion repressed ribosomal protein expression (Figure 2C). Because TORC1 inactivation increases both the chronological life span and resistance to heat stress in yeast,^43–45^ we tested whether the *aur1-1* mutant had similar effects. Indeed, we observed an extended lifespan and enhanced heat tolerance in *aur1-1* compared to *AUR1* (Figures 2D and 2E). Moreover, we tested whether *aur1-1* mutant cells had short telomeres, as TORC1 inactivation by rapamycin leads to telomere shortening.^46^ Northern blot analysis showed that telomeres in the *aur1-1* mutant were shorter than those in the *AUR1* strain (Figure 2F). Thus, these data indicate that sphingolipid-deficient cells display a phenotype similar to that observed under TORC1 inactivation conditions.

**Figure 2.**
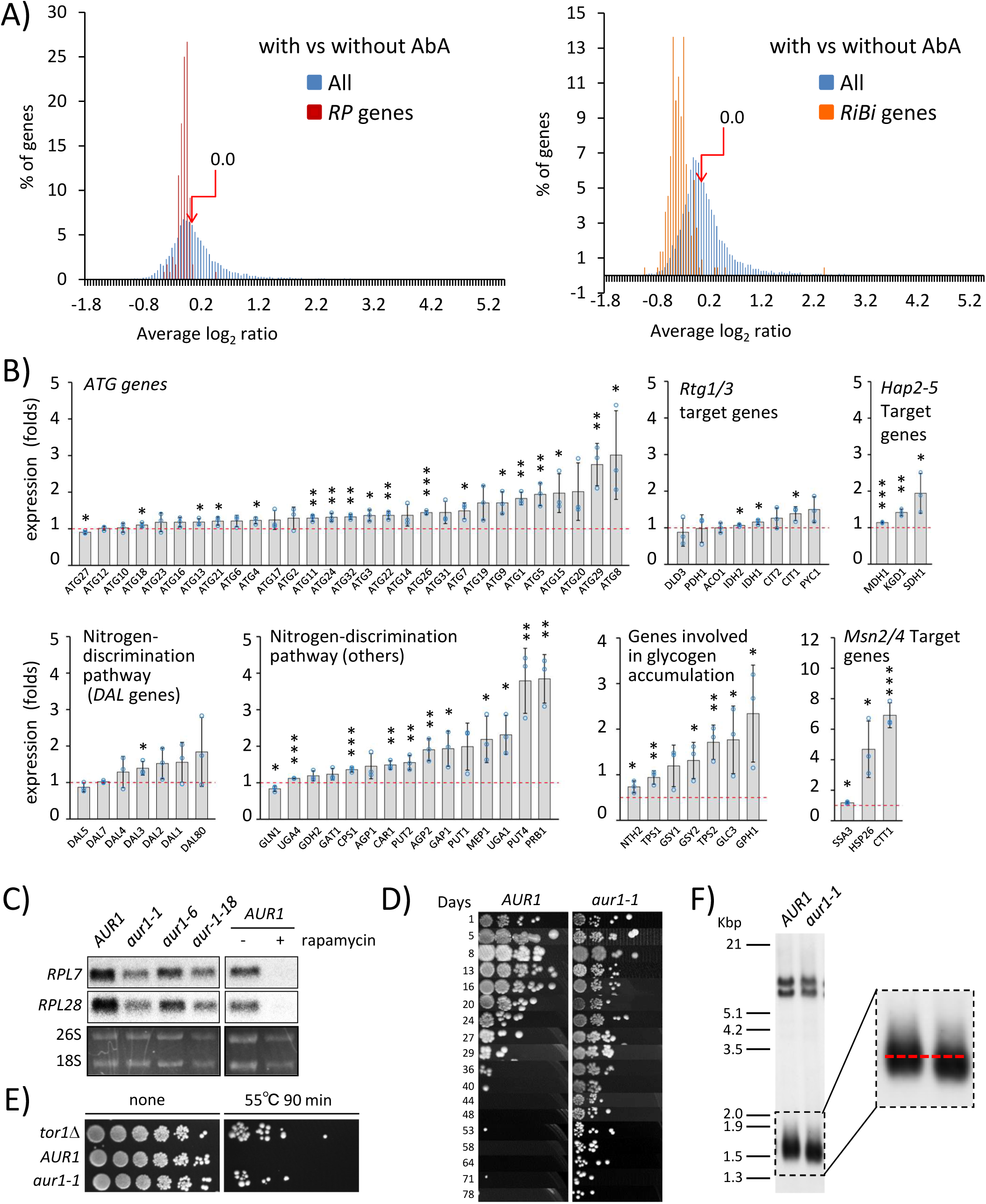
Sphingolipid-deficient cells display a set of phenotypes induced by reduced TORC1 activity. A, B) Microarray analysis of gene expression profiles of wild-type cells before and after treatment with 0.075 μg/ml AbA for 4h. The relative expression levels of 5541 genes before and after treatment with AbA were analyzed, and average values were obtained from three independent experiments. The results were expressed as percentage of ribosomal protein (RP) (120 genes) and all genes (A, left) or RiBi genes involved in ribosome biogenesis (220 genes) and all genes (A, right) with 0.05-fold difference in the relative expression levels. The relative fold expressions (the expression in cells treated with AbA relative to the expression in untreated cells) of *ATG* genes, Rtg1/3, Hap2-5, genes involved in the nitrogen-discrimination pathway and glycogen accumulation, and Msn2/4 target genes are shown (B). Average values and standard deviations (SD) were obtained from three independent experiments for each group. *P value for the Student’s t-Test < 0.05; **P < 0.01; ***P < 0.001. C) mRNA levels of ribosomal protein genes, *RPL7* and *RPL28,* in the indicated strains treated with or without 200 nM rapamycin for 45 min were determined by northern analysis. Ethidium bromide-stained rRNA was used as a loading control. D) Spot test chronological lifespan assay for *AUR1* and *aur1-1* mutant cells. The cells were grown in SC medium to the late stationary phase, and then aliquots from cultures at the indicated days were serially diluted tenfold, spotted onto YPD plates, and incubated for 4 days at 25°C. E) Heat stress resistance. Cells were cultured in SC medium for 3 days at 25°C and then exposed to heat shock (55°C for 90 min). Aliquots were serially diluted fivefold, spotted onto YPD plates, and incubated for 4 days at 25° °C. F) Telomere lengths of *AUR1* and *aur1-1* strains were analyzed by southern blot as described in the Method Details.

To directly test whether sphingolipids regulate TORC1 activity, we monitored cell sensitivity to rapamycin and phosphorylation of Sch9, the most sensitive readouts of TORC1 activation.^47^ We analyzed the effects of *aur1* mutations on rapamycin sensitivity using strains with BY and W303-genetic backgrounds to rule out strain-dependent effects. In both genetic backgrounds at the permissive temperature of 25L°C, the *aur1-1* and *aur1-18* mutations conferred high sensitivity to rapamycin, whereas the *aur1-6* mutation had little impact compared to the control strain (Figure 3A). In addition, *aur1-1* and *aur1-18* mutations conferred high sensitivity to caffeine (Figure S7A), another inhibitor of TORC1.^40^ Negative genetic interactions between *AUR1* and *TOR1* or *TCO89* (Figure 3B and S7B) also confirmed a functional link between sphingolipid metabolism and TORC1 signaling. As phosphorylation of Sch9 was decreased in *aur1* mutants and AbA-treated cells, similar to rapamycin-treated cells (Figure 3C and 3D), these results indicate that sphingolipids modulate TORC1 activation. Moreover, treatment with myriocin, a specific inhibitor of Lcb1/Lcb2^48^, and the *lcb1-100* mutation reduced Sch9 phosphorylation (Figure 3D and 3E). The *lcb1-100* mutant also exhibited hypersensitivity to rapamycin (Figure 3F), induced autophagy (Figure 3G), and repressed ribosomal protein expression (Figure 3H). These data, together with the evidence that the *LCB1* gene exhibits a negative genetic interaction with the *TOR1* gene (Figure 3E, 3F, and 3G), suggest that decreased levels of complex sphingolipids, but not increased ceramide levels, contribute to TORC1 inhibition. As *csg1*Δ*csg2*Δ and *ipt1*Δ were not hypersensitive to rapamycin but showed stronger resistance than wild-type cells (Figure 3I), this implies that the reduction in IPC levels is the main reason for the reduced TORC1 activity in the *aur1* mutant strain.

**Figure 3.**
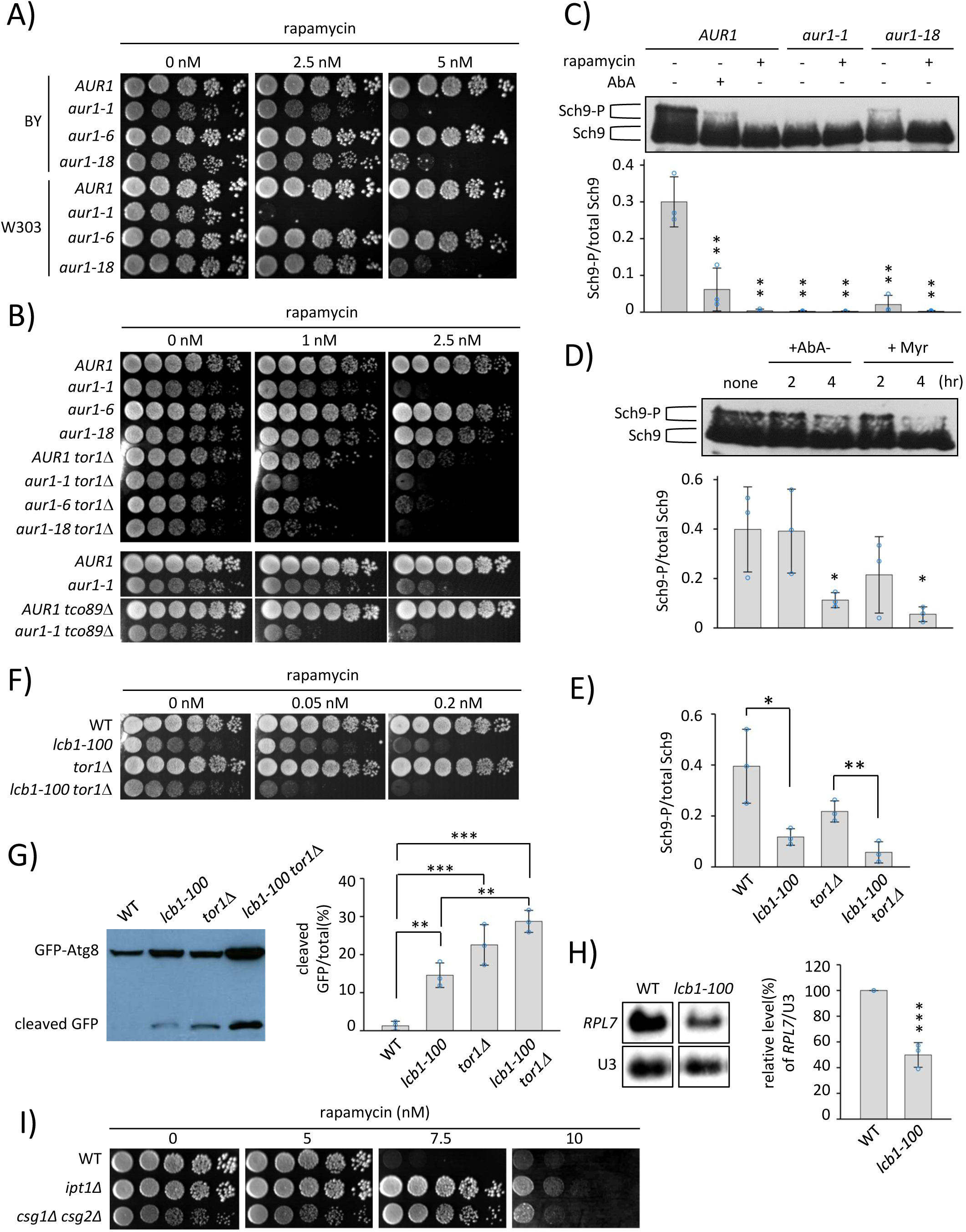
Inositolphosphorylceramide (IPC) is required for TORC1 activation. A, B, F, and I) Rapamycin sensitivity. Cells were serially diluted fivefold, spotted onto YPD plates containing the indicated concentrations of rapamycin, and incubated for 4 days at 25°C. C, D, and E) Sch9 phosphorylation. Cells expressing Sch9-5HA were grown in YPD and treated with or without 200 nM rapamycin for 30 min (C), 0.5 μg/ml AbA for 2h (D) or 4 h (C, D), 200 μg/ml myriocin (Myr) for 2h or 4h (D) at 25°C. NTCB-treated protein extracts were analyzed using western blotting. Sch9 phosphorylation was assessed as the ratio of phosphorylated (Sch9-P)/total Sch9 and expressed as the mean ± SD from three independent experiments (C, D, E). *P value for the Student’s t-Test < 0.05; **P < 0.01. G) Cells expressing GFP-Atg8 were grown exponentially in YPD and analyzed by western blotting (left), and autophagic flux was expressed as the mean ± SD from three independent experiments (right). **P value for the Tukey-Kramer multiple comparison test < 0.01; ***P < 0.001. H) Levels of *RPL7* mRNA and U3 snoRNA (as a loading control) were analyzed by northern blotting (left), quantified from three independent experiments, normalized to U3, and expressed as the mean ± SD of relative levels (right). ***P value for the Student’s t-Test, < 0.001.

### Sphingolipids are required for EGO complex-mediated TORC1 activation via vacuole Lo domain formation

Previous studies have shown that TORC1 activation is mediated via the EGO complex consisting of Ego1, Ego3, Grt1, and Gtr2, which reside on the vacuole by anchoring to the membrane through the N-terminal lipid modification of Ego1.^5,49^ We therefore tested if the EGO complex is involved in sphingolipid-dependent regulation of TORC1 activation. First, we examined the effects of Gtr1 and Gtr2 overexpression on rapamycin hypersensitivity in *aur1* mutants. As shown in Figure 4A, overexpression of Gtr1 or Gtr2 partially rescued the sensitivity of *aur1* mutants to rapamycin. We then analyzed whether the *aur1* mutation affected the localization of the EGO complex (Figure 4B). Fluorescence images and subsequent quantitative analysis indicated that Gtr1-GFP expressed under the control of the endogenous *GTR1* promoter was localized to the vacuole membrane in *AUR1* or wild-type cells. However, consistent with a previous observation,^50^ the localization of Gtr1-GFP in vacuoles was significantly reduced in *ego3*Δ mutant cells, with the majority of Gtr1-GFP being mislocalized to the cytoplasm. This mislocalization of Gtr1-GFP was also observed in *aur1-1* mutant cells. In contrast, the Ego1-GFP signal in the *aur1-1* mutant remained on the vacuolar membrane, as in the control *AUR1* strain. The finding that sphingolipids regulate the localization of Gtr1-GFP led us to consider two possible models for sphingolipid-dependent regulation of TORC1 activation: (1) Gtr1 mislocalization is directly responsible for TORC1 inhibition in sphingolipid-deficient mutants, or (2) Gtr1 mislocalization, together with additional factors, cooperatively represses TORC1 activity. To clarify which model was correct, we investigated whether artificial tethering of Gtr1 to the vacuole membrane could rescue rapamycin sensitivity in *aur1-1* or *ego3*Δ mutant cells. We utilized previously established strains expressing Ego1-GFP to form an artificial tether, as Ego1-GFP remains on the vacuolar membrane in the mutants and can interact with Gtr1 fused to a C-terminal GFP-binding protein (Gtr1-GBP). We observed that artificial tethering via GFP-GBP interaction suppressed rapamycin sensitivity in the *ego3*Δ strain, which is similar to previously reported results,^49^ but not in the *aur1-1* mutant (Figure 4C). We also confirmed that the mislocalization of Gtr1-GFP in the *aur1-1* mutant was rescued by artificial tethering through the interaction of GFP and GBP (Figure S8A); however, the sensitivity to rapamycin remained high (Figure S8B). These data indicate that vacuolar-tethered Gtr1-GFP in the *aur1-1* mutant is not sufficient to promote TORC1 activity, and factors other than Gtr1 mislocalization are likely to contribute to the inhibition of TORC1 activity in the *aur1* mutant.

**Figure. 4.**
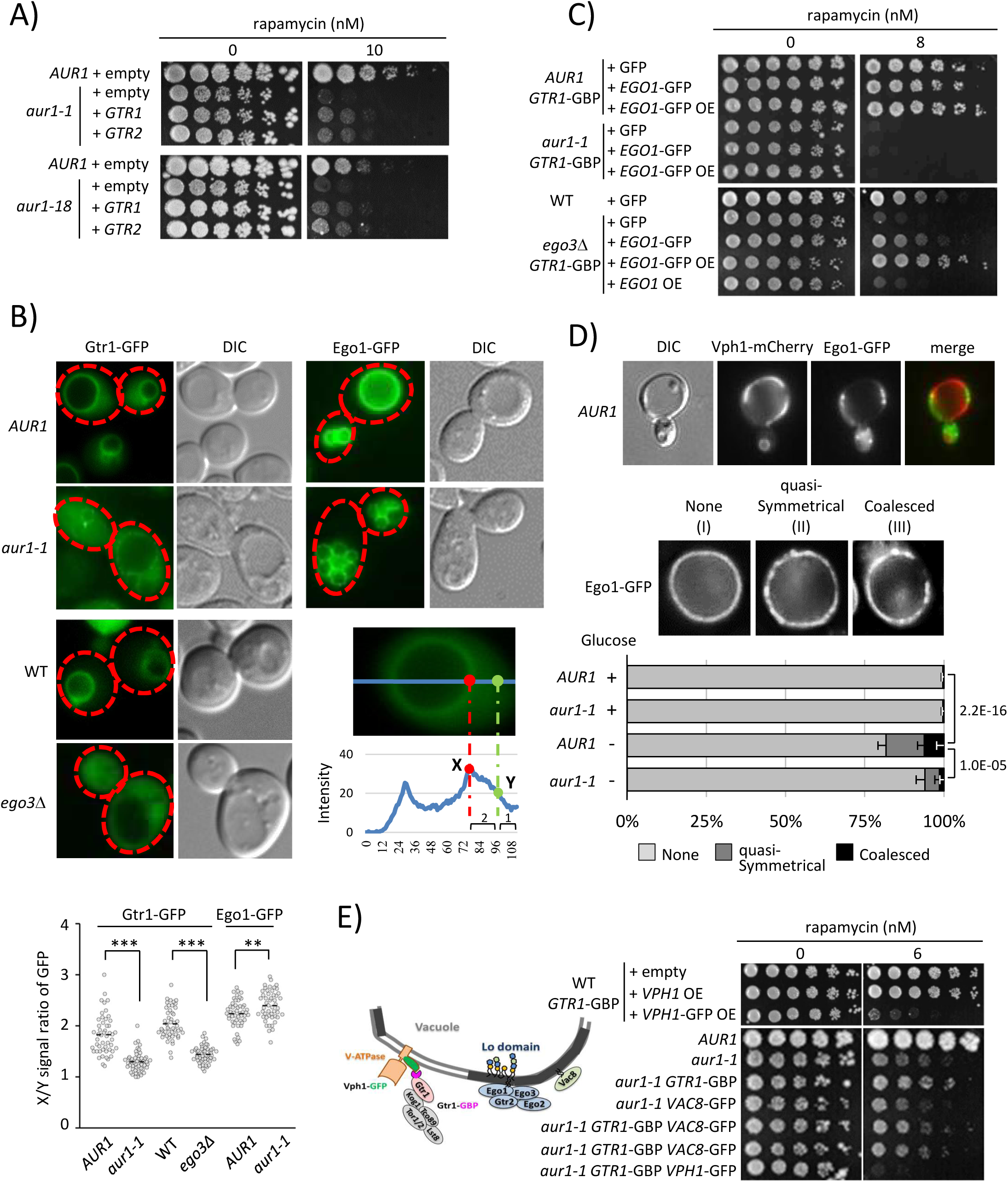
Sphingolipid-enriched vacuole liquid-ordered (Lo) domains are required for EGO complex-mediated TORC1 activation. A, C, and E) Rapamycin sensitivity. Cells were serially diluted fivefold, spotted onto YPD plates containing the indicated concentrations of rapamycin, and incubated for 4 days at 25°C. Schematic representation of GFP-GBP fusion protein interaction-mediated anchoring of Gtr1 to Vph1 in the vacuolar Ld domain (left panel in E). B) Cells expressing Gtr1-GFP or Ego1-GFP were grown at 25°C in SD medium and observed using fluorescence microscopy. The right panel in the middle is an intensity plot along the blue line in wild-type cells, and X and Y indicate the positions of the vacuolar membrane and cytosol (located in the two-thirds region between the vacuolar membrane and the plasma membrane), respectively. X/Y intensity ratios were analyzed in three independent experiments, with a minimum of 50 cells counted per genotype, and the data are represented as beeswarm plots (left panel at the bottom of B). **P value for the Student’s t-Test < 0.01; ***P < 0.001. D) Cells expressing Vph1-mCherry and Ego1-GFP were grown at 25 °C in SD medium, washed, resuspended in SD (Glucose (+)) or SD lacking glucose (Glucose (-)), and incubated at 25 °C for 24 h. Cells were visualized using fluorescence microscopy. Photographs obtained from *AUR1* cells in SD lacking glucose are shown, and quasi-symmetrical and coalesced microdomains are shown in (II) and (III), respectively. The percentage of cells with vacuolar domains was calculated from 100 cells for each sample per experiment and expressed as the mean ± SD from three independent experiments. Statistical significance was assessed using Fisher’s exact test.

To clarify and gain further insights into the mechanism underlying sphingolipid/EGO complex-dependent regulation of TORC1 activation, we measured the amino acid content of whole cells and vacuolar pools in *AUR1* and *aur1* mutant cells. This is due to the fact that EGO complex-mediated activation of TORC1 depends on the amino acid availability^51^, and has been linked to sphingolipid metabolism.^52–54^ In both pools, contents of arginine were particularly reduced in *aur1* mutants (Figure S9A and S9B), raising the possibility that the reduced arginine level is an important contributor to TORC1 inhibition in *aur1* mutant cells. However, this was not the case, as *CAN1* deletion, which significantly reduced arginine availability (Figure S9A and S9B), did not affect the rapamycin sensitivity of control *AUR1* and *aur1* mutant cells (Figure S9C).

The EGO complex and TORC1 have been shown to localize to vacuole Lo membrane domains,^10,12,13,15,17^ which are phase-separated from the Ld domains marked by the vacuolar ATPase subunit Vph1 during glucose starvation (Figure 4D). The vacuole Lo domains are sterol-enriched, wherein sterols are necessary for domain formation.^12^ A recent study showed that cells with reduced *AUR1* expression predominantly contain vacuoles lacking domains in the early stationary stage, as judged using the Ld vacuolar domain marker, Pho8-GFP,^14^ suggesting a role of sphingolipids in Lo domain formation. Therefore, we examined whether the *aur1* mutation affects Lo domain formation in vacuolar membranes. As shown in Figure 4D, *aur1-1* mutant cells had fewer visible domains during glucose starvation than control *AUR1* cells, confirming that sphingolipids are required for Lo domain formation. These results suggest that vacuolar Lo domains may modulate TORC1 activation. To test this, we introduced an artificial tether of Gtr1-GBP with Vph1-GFP into the cells and assessed whether the mislocalization of Gtr1 to Ld domains affected rapamycin sensitivity (Figure 4E). Artificial tethering of Gtr1 (Gtr1-GBP) to Ld domains (Vph1-GFP) increased the sensitivity of both wild-type and *aur1-1* cells to rapamycin. In contrast, as a negative control, we did not observe a significant increase in rapamycin sensitivity in the *aur1-1* mutant expressing Gtr1-GBP alongside Vac8-GFP, which is targeted to vacuolar Lo domains via its N-terminal lipid modification sites.^55^ These findings strongly imply that vacuolar Lo domains are required to maintain sufficient and functional EGO complex-mediated TORC1 activity.

To validate the involvement of sphingolipid-dependent Lo domains in TORC1 activation, we next used an engineered strain, *GhLag1,* that produces cellular membranes with shorter ceramides (C16-C18 ceramides) than the wild-type strain (C26 ceramides).^56,57^ Thus, this *GhLag1* mutant generates shorter IPC species than the wild-type cells.^58^ Although biophysical studies using model membranes showed that both very long (C24) and short (C16-C18) acyl chain ceramides induce phase separation, it is expected that shorter-chained sphingolipids in cell membranes would have different biophysical impacts on domain biogenesis and maintenance, since shorter sphingolipids are not able to form interdigitated phases.^59^ Indeed, we found that the *GhLag1* cells have a negative impact on vacuole Lo domain formation (Figure 5A). To learn more about the biophysical characteristics of biomembranes composed of short sphingolipids, we extracted protein-free lipids from wild-type and *GhLag1* cells, generated bilayers in vitro, and assessed the surface topography of these membranes using atomic force microscopy (AFM). The AFM images revealed that the wild-type and *GhLag1* membranes exhibited lateral lipid phase segregation with lipid domains of different heights: 8 nm and 4 nm in the wild-type and *GhLag1*, respectively (Figure 5B, upper and middle panels), suggesting that the Lo domains in the wild-type and *GhLag1* membranes have different thicknesses. Moreover, AFM force spectroscopy revealed that the breakthrough force (force required to pierce through the bilayer) values in the *Ghlag1* membranes were lower than those in the wild-type membranes (Figure 5B, lower panel). This suggests that *GhLag1* membranes have a lower membrane order than wild-type membranes. Consistently, liposomes composed of lipids extracted from *GhLag1* cells exhibited lower generalized polarization (Figure 5C) and lifetime (Figure 5D) of the solvatochromic probe laurdan, which are indicators of increased membrane fluidity.^60^ Similarly, a low generalized polarization has also been observed in model membranes made from lipid extracts of deletion mutant *ELO3*, the gene encoding a fatty acid elongase, which is unable to synthesize very long chain sphingolipids (C26).^61^ Finally, *Ghlag1* cells exhibited high sensitivity to rapamycin (Figure 5E), reduced localization of Gtr1-GFP to the vacuolar membrane (Figure 5F), and decreased Sch9 phosphorylation (Figure 5G). Collectively, these findings support the model that sphingolipid-mediated vacuolar Lo domain formation is necessary for TORC1 activation.

**Figure. 5.**
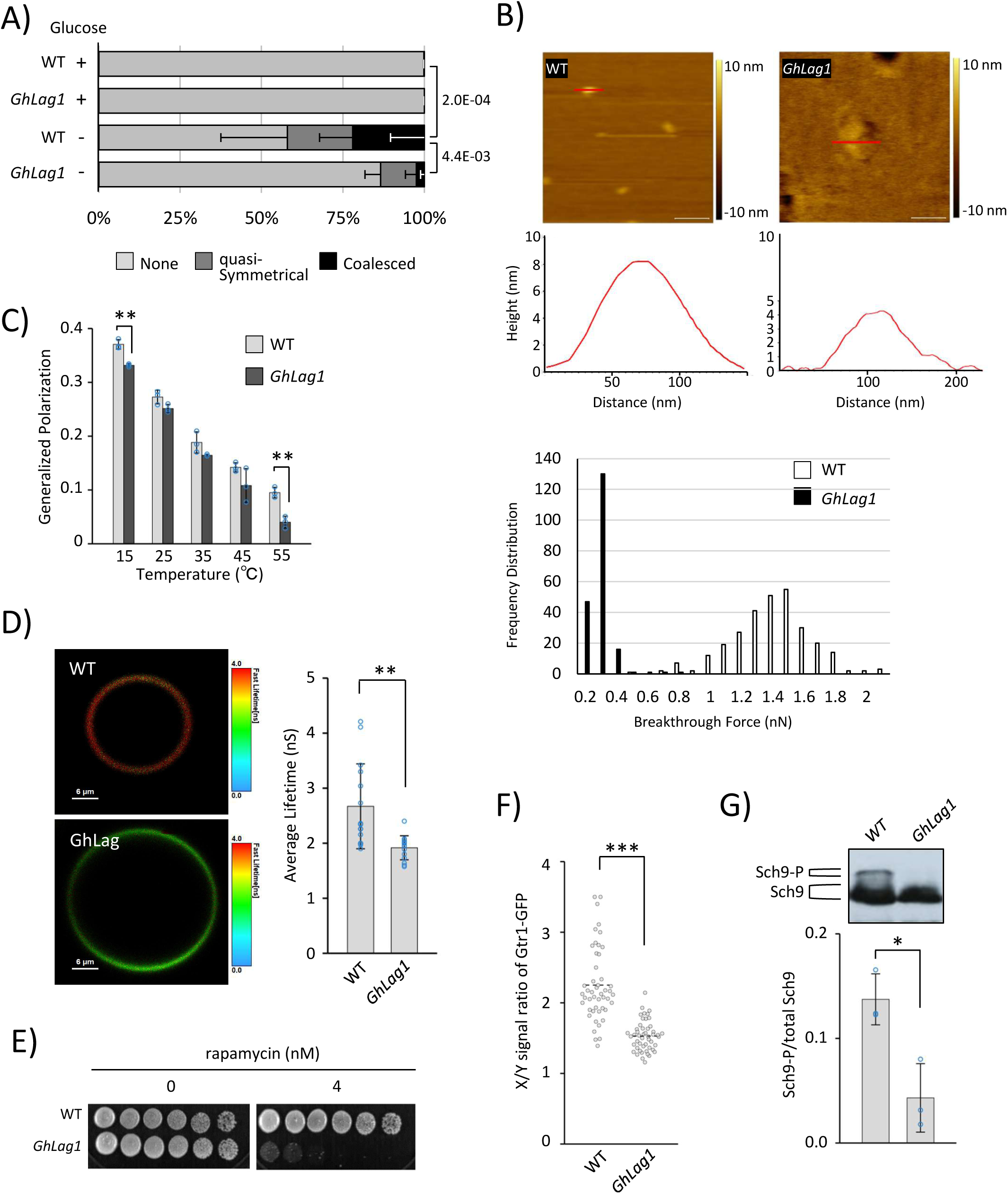
Chain length of sphingolipids is critical for Lo domain formation and TORC1 activation. A) Wild-type and *GhLag1* cells expressing Ego1-GFP were incubated in SD or SD lacking glucose at 25 °C for 24 h and visualized by fluorescence microscopy as described in Fig 4D. The percentage of cells with the vacuolar domains was calculated from 100 cells for each sample per experiment and expressed as the mean ± SD from three independent experiments. Statistical significance was tested using Fisher’s exact test. B) 2D Topographical AFM images of lipids from wild-type and *GhLag1* (upper panel) (Scale bar: 200 nm), height profile of the selected region marked in red (middle panel), and breakthrough force distribution (lower panel). For wild-type, the height difference between liquid-ordered (Lo) and -disordered (Ld) phases is ∼8 nm, whereas for *GhLag1*, the height difference is ∼4 nm, indicating that Lo/Ld phase height difference in *GhLag1* membrane is smaller than that in wild-type membrane. The mean values of the breakthrough force distribution were 1.39 nN ± 0.25 nN (n=287) and 0.29 nN ± 0.06 nN (n=197) for wild-type and *GhLag1*, respectively, showing differences in nanomechanical stability in the two lipid systems. C) Temperature-dependent Laurdan generalized polarization in LUVs made with lipids from wild-type and *GhLag1*. Data are presented as mean ± SD of three independent experiments. Student’s t test: **p ≤ 0.01. D) Representative FLIM images (left panel) and average fluorescence lifetime (right panel) of Laurdan in GUVs made with lipids from wild-type and *GhLag1*. The lifetime bar graph was constructed using at least 15 independent images from each lipid system. Student’s t test: **p ≤ 0.01. E) Rapamycin sensitivity. Cells were serially diluted fivefold, spotted onto YPD plates containing the indicated concentrations of rapamycin, and incubated for 4 days at 25 °C. F) Gtr1-GFP localization in wild-type and *GhLag1*. The X/Y intensity ratios of Gtr1-GFP were quantified, and the data are represented with beeswarm plots, as described in Fig. 4B. Student’s t-test: ***p ≤ 0.001. G) Sch9 phosphorylation. NTCB-treated protein extracts from wild-type and *GhLag1* cells expressing Sch9-5HA were analyzed by western blotting, and the ratio of phosphorylated (Sch9-P)/total Sch9 was quantified and expressed as the mean ± SD from three independent experiments, as described in Fig. 3C. Student’s t test: *p ≤ 0.05.

### Sphingolipids integrate TORC2 with TORC1 through vacuolar Lo domain formation

Given our present results and the fact that TORC2 signaling upregulates sphingolipid synthesis,^62^ we hypothesized that TORC2 signaling modulates TORC1 activation through sphingolipid metabolism. Temperature-sensitive mutants of TORC2 components or proteins that function downstream or upstream of TORC2 (Figure S10A)^63^ were used to test this hypothesis. Under permissive temperature conditions, mutations in Avo3, one of the TORC2 components, as well as its upstream regulators Slm1/Slm2, Stt4, Mss4, and downstream effectors Ypk1/Ypk2, caused hypersensitivity to rapamycin (Figure 6A and S10B). A double mutant of Pkh1/Pkh2, critical upstream activators of Ypk1/Ypk2, was also rapamycin hypersensitive, whereas mutations of Pkc1 and Rho1, upstream activators of Pkc1, did not alter rapamycin sensitivity. The mutants that exhibited high sensitivity to rapamycin had reduced expression of ribosomal proteins (Figure 6B and S10C) and reduced vacuolar localization of Gtr1 (Figure 6C and S10D). Moreover, as additional indicators of TORC1 inactivation, we observed a reduction in Sch9 phosphorylation (Figure 6D) and induction of autophagy (Figure 6E) in the *avo3-30* mutant cells. Hypersensitivity to rapamycin was also observed using the *ypk1ts ypk2*Δ strain, and the same was rescued by the expression of wild-type Ypk1, but not by the expression of *ypk1* mutants *YPK1* S644A/T662A and *YPK1* T504A, which carry mutations in the TORC2 and Pkh1/2 phosphorylation sites,^64,65^ respectively (Figure S10E). This result suggests that Ypk1 lies downstream of both TORC2 and Pkh1/2 in the TORC1 activation cascade. This is further supported by our observation that the expression of a constitutively active allele of Ypk1 (*YPK1* D242A)^64,65^ rescued the rapamycin sensitivity of *avo3-30* cells (Figure 6F) and *slm1-1 slm2*Δ cells (Figure S10F). Additionally, expression of *YPK1* D242A restored autophagy induction (Figure 6G) and Gtr1 mislocalization (Figure 6H) in *avo3-30* cells. We also showed that the *avo3-30* mutation reduced Lo domain formation during glucose starvation and that this phenotype was rescued by the expression of *YPK1* D242A (Figure 6I). These results suggest that Ypk1/2 acts downstream of TORC2 to regulate TORC1 activation via Lo domain formation.

**Figure 6.**
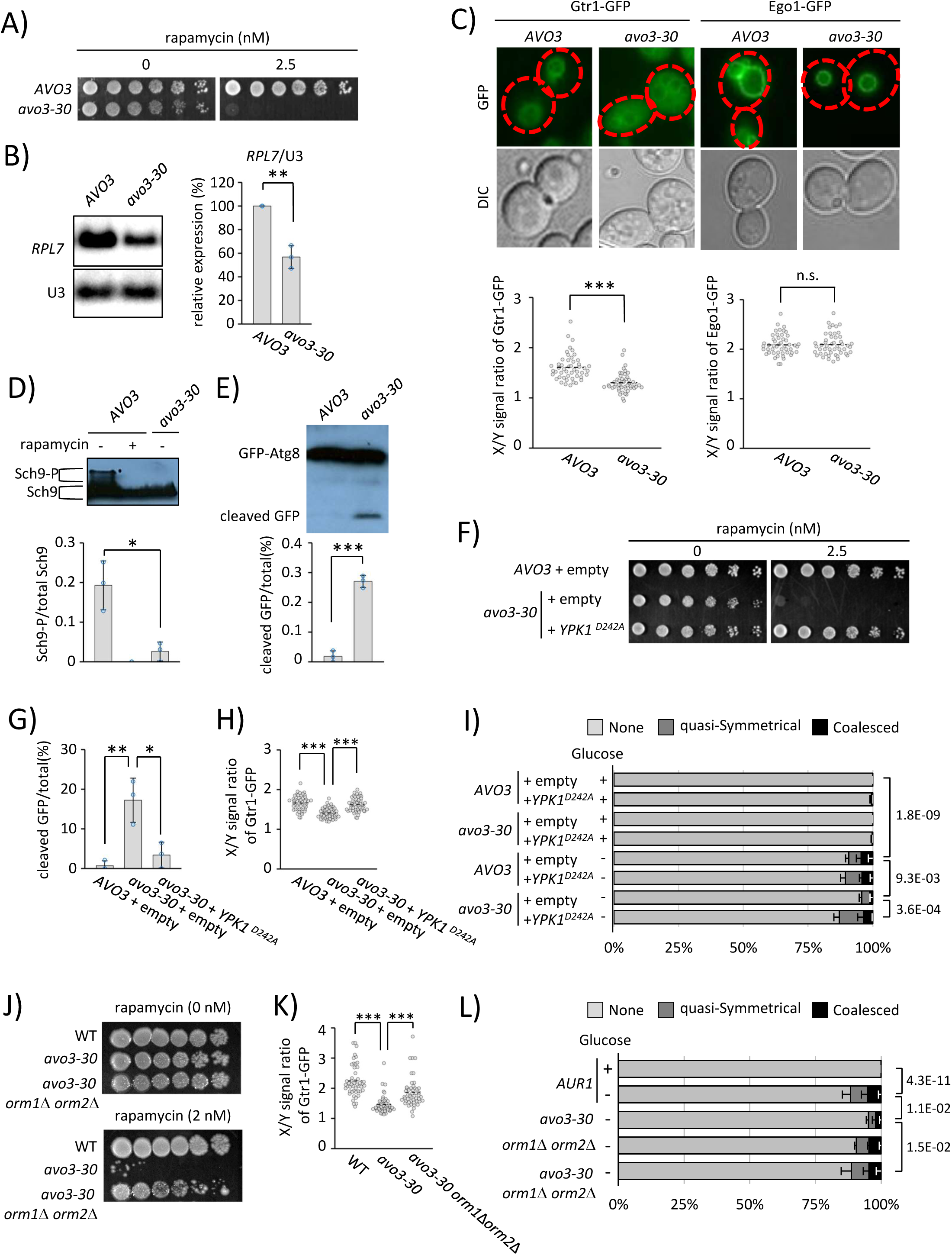
TORC1 is functionally linked to TORC2 via the Ypk/Orm regulatory pathway of sphingolipid metabolism. A, F, and J) Rapamycin sensitivity of *AVO3* and mutants was determined by spotting serial dilutions of cells onto YPD plates with or without the indicated concentrations of rapamycin (A, J). *AVO3* and mutant cells transformed with empty plasmid or plasmids expressing Ypk1D242A (F) were spotted onto SC plates lacking uracil and methionine (SC-UM) with or without 2.5 nM rapamycin. The cells were then grown for 4 days at 25 °C. B) Levels of *RPL7* mRNA and U3 snoRNA were analyzed by northern blotting (left panel), quantified from three independent experiments, normalized to U3, and expressed as the mean ± SD of relative levels (right panel), as described in Fig. 3H. Student’s t test: **p ≤ 0.01. C, H, and K) Cells expressing Gtr1-GFP or Ego1-GFP transformed without (C, K) or with empty plasmid or plasmid expressing Ypk1D242A (H) were observed by fluorescence microscopy. X/Y intensity ratios were quantified, and the data are represented with beeswarm plots, as described in Fig. 4B. ***P value for the Student’s t-Test, < 0.001 (C), Tukey-Kramer multiple comparison test (H, K). D) Phosphorylation of Sch9 was monitored by western blotting (D, left panel), and the ratio of phosphorylated (Sch9-P)/total Sch9 was expressed as the mean ± SD from three independent experiments (D, right panel), as described in Fig. 3C. Student’s t test: *p ≤ 0.05. E, G) Autophagic degradation of GFP-Atg8 was analyzed by western blotting (E, left panel), and autophagic flux was expressed as the mean ± SD from three independent experiments (E, right panel, and G). Student’s t-test: ***p ≤ 0.001 (E). Tukey-Kramer multiple comparison test: **p ≤ 0.01; *p ≤ 0.05 (G). I, L) Cells expressing Ego1-GFP transformed with empty plasmid or plasmid expressing Ypk1D242A (I) or without (L) were incubated in SD lacking uracil and methionine (SD-UM) or SD-UM lacking glucose (I) and in SD or SD lacking glucose (L) at 25°C for 24 h, as described in Fig 4D. The percentage of cells with vacuolar domains was calculated from 100 cells and expressed as the mean ± SD of three independent experiments. Statistical significance was tested using Fisher’s exact test.

TORC2-Ypk1/2 signaling phosphorylates the regulatory proteins Orm1 and Orm2 to cancel the repression of long-chain base (LCB) synthesis, which is the first step of *de novo* sphingolipid synthesis.^63,66^ Therefore, we sought to determine whether Orm1/2 also influences TORC1 activity in *avo3-30* cells. We observed that double deletion of *ORM1* and *ORM2* rescued rapamycin hypersensitivity in *the avo3-30* mutant (Figure 6J). In addition, the mislocalization of Gtr1-GFP (Figure 6K) and reduced Lo domain formation (Figure 6L) in *avo3-30* cells were suppressed by *ORM1*/*ORM2* deletion. Consistent with these results, treatment of *avo3-30* cells with phytosphingosine (PHS), a major LCB in yeast, significantly rescued rapamycin sensitivity (Figure S10G) and Gtr1 mislocalization (Figure S10H) in *avo3-30* cells, indicating that sphingolipids are involved in TORC2-Ypk1/2-dependent regulation of TORC1 activation. As a control, treatment of *lcb1-100* cells with PHS rescued rapamycin sensitivity (Figure S10I), whereas the expression of *YPK1* D242A did not rescue rapamycin sensitivity in *lcb1-100* or *aur1-1* cells (Figure S10F). This result confirmed that sphingolipids function downstream of Ypk1 to activate TORC1.

Ypk1 has been shown to positively regulate ceramide synthesis by directing the phosphorylation of ceramide synthases Lac1 and Lag1, and calcineurin antagonizes this Ypk1-dependent phosphorylation.^67^ Therefore, we tested whether calcineurin plays an antagonistic role in the TORC2-Ypk1/2-dependent regulation of TORC1 activation. This was the case; deletion of *CNB1*, which encodes the calcineurin regulatory subunit B, partially restored the rapamycin sensitivity (Figure S11A), reduced phosphorylation of the direct TORC1 substrate, Rps6 (Figure S11B), and reduced Lo domain formation (Figure S11C), in *avo3-30* cells. Taken together, these results support the model that sphingolipids integrate TORC2 with TORC1 through vacuolar Lo domain formation.

### Vacuole fusion and acidification regulate TORC1 activation through IPC synthesis

Given that genes involved in vacuolar fusion and acidification exhibit negative genetic interactions with *AUR1* (Figure 1A, S1, S2, and S3), we postulated that deletion of these genes might influence TORC1 activity. Accordingly, we analyzed rapamycin sensitivity and Sch9 phosphorylation in the deletion mutants. As expected, and as previously reported,^68–71^ deletion of genes such as *VPS16* and *VPS11*, which encode components of the Vps-C core required for vacuole membrane fusion, conferred high sensitivity to rapamycin (Figure 7A) and decreased Sch9 phosphorylation (Figure 7B), confirming the reduced TORC1 activity in the mutants. Consistent with this, we observed Gtr1 mislocalization in Vps-C core mutants (Figure 7C). We then measured complex sphingolipid synthesis and found that the production of IPC-C, the main IPC in *Saccharomyces cerevisiae,* which contains the PHS-C26OH ceramide, as well as MIPC, was significantly reduced in the Vps-C core mutants (Figure 7D and 7E). Similar positive correlations between reduced IPC levels and increased rapamycin sensitivity (as well as increased AbA sensitivity (Figure S2B)) or reduced Sch9 phosphorylation were observed in vacuole acidification-defective mutants, such as *vma6*Δ and *vma22*Δ (Figure S12). These results reveal that vacuolar fusion and acidification play essential roles in the sphingolipid-dependent regulation of TORC1 activation. However, the expression of *YPK1* D242A did not suppress rapamycin sensitivity in *vps16*Δ and *vma6*Δ, vacuolar fusion- and acidification-defective mutants, respectively (Figure 7F), indicating that the regulation of TORC1 by vacuolar fusion and acidification is not linked to TORC2-Ypk1/2 signaling.

**Figure 7.**
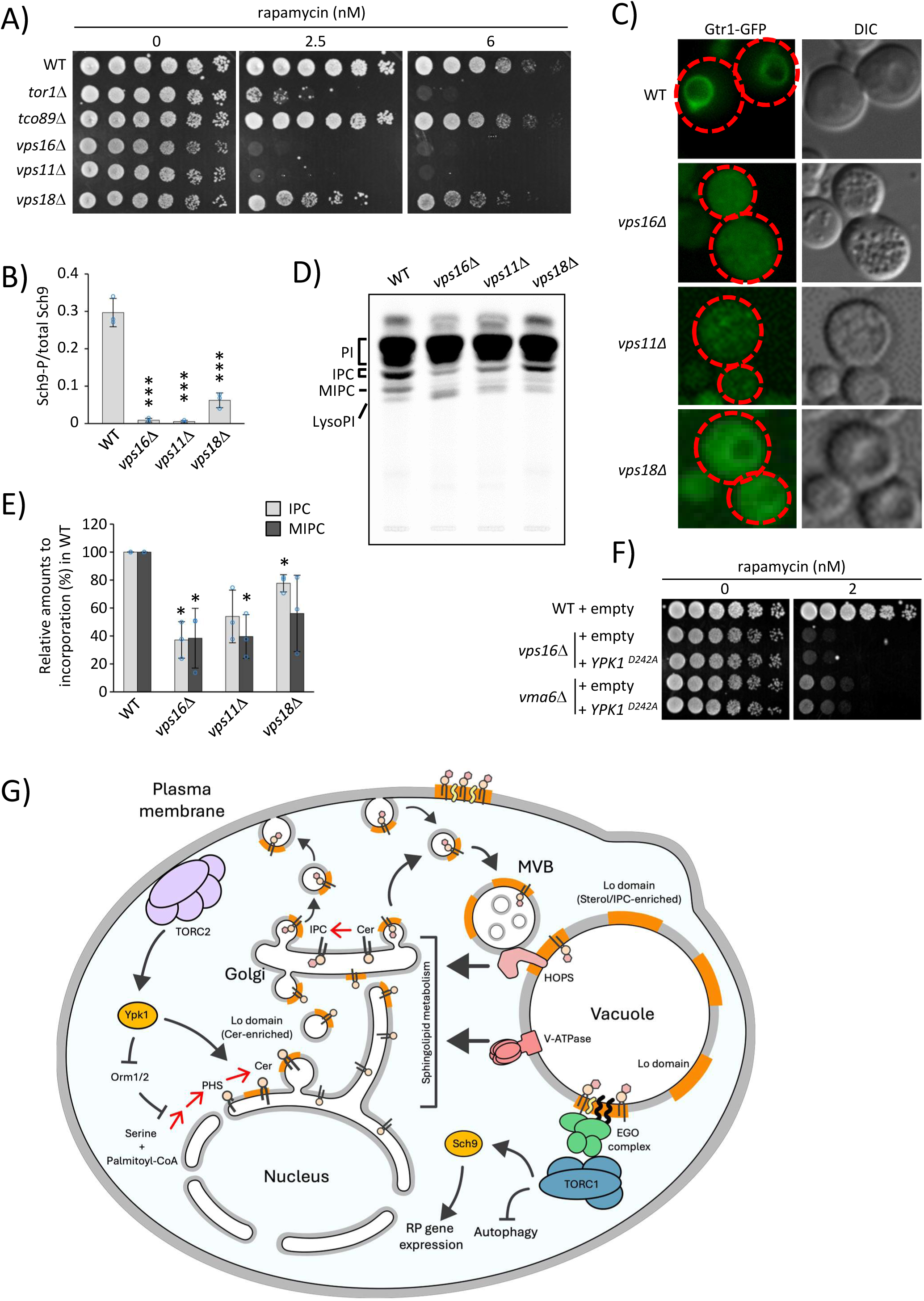
The Vps-C core regulates TORC1 activation via IPC synthesis. A, F) Rapamycin sensitivity of wild-type and mutants was determined by spotting serial dilutions of cells onto YPD plates with or without the indicated concentrations of rapamycin (A). Wild-type and mutant cells transformed with empty plasmid or plasmid expressing Ypk1D242A were spotted onto SC plates lacking uracil and methionine (SC-UM) with or without 2 nM rapamycin (F). The cells were then grown for 4 days at 25 °C. B) Phosphorylation of Sch9 was analyzed, and the ratio of phosphorylated (Sch9-P)/total Sch9 was expressed as the mean ± SD from three independent experiments. Student’s t-test: ***p ≤ 0.001. C) Gtr1-GFP localization in wild-type and Vps-C core mutants was analyzed using fluorescence microscopy. D, E) Cells were labeled with [^3^H]*myo*-inositol for 90 min at 25 °C. The labeled lipids were analyzed by thin-layer chromatography (TLC) using solvent system I (D), and the relative amounts of incorporation (%) into IPC or MIPC in wild-type cells were determined from three independent experiments and expressed as the mean ± SD (E). Student’s t test: *p ≤ 0.05. G) Model of functional linkage between TORC1 and TORC2 via sphingolipid metabolism and vacuolar Lo domain. TORC2 and VPS-C core (CORVET/HOPS)/V-ATPase regulate sphingolipid metabolism via Ypk/Orm-dependent and -independent pathways, respectively. In vacuoles, the sphingolipid-enriched membrane Lo domain is required for EGO complex-mediated TORC1 activation. The reduction of complex sphingolipids, most likely IPC, triggers vacuolar fragmentation, probably by disturbing vacuole-vacuole fusion.

## DISCUSSION

Over the last few decades, researchers have introduced the concept of lipid rafts, proposing that lipid-lipid interactions laterally organize biological membranes into domains with different structures, compositions, and functions. This concept has provided insights and developments across many disciplines related to membrane biology. However, the existence and functional relevance of these domains in cells remain unclear. In this study, we discovered an essential role for the Lo lipid domains of the vacuolar membrane, formed by sphingolipids, in the activation of TORC1 in budding yeast. Importantly, TORC2-Ypk1/2 signaling, which modulates sphingolipid synthesis, controls TORC1 activation through the formation of Lo domains. Thus, we believe that sphingolipids integrate TORC2 and TORC1 via vacuolar Lo domains.

Our data demonstrate that TORC1 activity is significantly reduced in mutant strains defective in sphingolipid synthesis, which is required for Lo domain formation, suggesting a critical role of the Lo domain in the activation of TORC1. Notably, our experiments with engineered strains that allowed artificial anchoring of Gtr1 to the Ld domain-localized Vph1 or the Lo-domain localized Ego1 and Vac8 pinpointed a functional link between the Lo domain and TORC1 signaling. This is in line with the observation that C16-C18 short acyl chain sphingolipids, which lower the degree of lateral lipid order, impair TORC1 signaling. As short sphingolipids cannot form interdigitated phases,^59^ it is conceivable that specific, stable domains in the cytoplasmic leaflet organized through interdigitation with very long acyl chain (C26) sphingolipids in the vacuolar luminal leaflet are essential for TORC1 activation. Intriguingly, *csg1*Δ*csg2*Δ and *ipt1*Δ cells did not show high sensitivity to rapamycin, suggesting that IPC is sufficient for the formation of interdigitated Lo domains to activate the TORC1. Furthermore, our finding that *csg1*Δ*csg2*Δ and *ipt1*Δ strains are resistant to rapamycin supports the idea that IPC accumulation due to impaired conversion to mannosylated IPC promotes TORC1 activity. However, the mechanism by which IPC-enriched Lo domains affect TORC1 activation remains unknown. As Vph1-free vacuolar Lo domains that form upon glucose starvation contain sterols,^12^ sphingolipid- and Ego1-containing Lo domains also likely contain sterols. Consistently, we found that treating cells with fenpropimorph, an ergosterol biosynthesis inhibitor that has previously been described to reduce vacuolar domain formation,^12^ inhibits the phosphorylation of Rps6 (Figure S13A) and induces autophagy even when nitrogen sources are abundant (Figure S13B). It should be noted that in nutrient-rich media, active TORC1 localizes to the vacuolar membrane, and Gtr1-GFP marked vacuoles appear as smooth rings lacking large coalesced domains. In contrast, clear domain segregation becomes apparent upon glucose starvation or endoplasmic reticulum stress. However, we cannot exclude the possibility that vacuolar domains are formed under nutrient-rich conditions but remain too small to be resolved by fluorescence microscopy. Interestingly, although our results demonstrate that sphingolipids are essential for Gtr1-GFP recruitment to the vacuole and for TORC1 activity, artificially tethering Gtr1-GFP to the vacuole is insufficient to restore TORC1 activity, suggesting the involvement of additional sphingolipid-dependent factors. Putative nanoscale Lo domains formed under nutrient-rich conditions may contribute to tethering TORC1 to the vacuole via interactions with the EGO complex and regulate the function of the EGO complex itself. Because the nucleotide-binding state of Gtr1-Gtr2 is crucial for the control of TORC1^72^, these Lo domains may affect this state. Moreover, the observation that anchoring Gtr1 to Ego1, which is assembled on the cytoplasmic leaflet side of the Lo domain, can rescue the growth defect of *ego3*Δ cells, implies that Ego3 functions primarily to recruit Gtr1 to the EGO complex rather than to directly modulate Gtr1 activity. In contrast, EGO complex-positive large coalesced domains, which can be visualized under glucose starvation, may be formed to inhibit TORC1 activity. The latter hypothesis has been proposed in a previous study.^10^

The integration of TORC1 with TORC2 via the vacuolar/lysosomal Lo domain is likely conserved from yeasts to mammals. Several reasons support this. First, the regulation of sphingolipid synthesis by TORC2 appears to be conserved in yeast and mammals. Yeast TORC2 promotes sphingolipid synthesis via post-translational regulation of serine palmitoyltransferase and ceramide synthase.^63^ Although it remains to be determined whether mammalian mTORC2 can similarly activate sphingolipid synthesis via post-translational regulation of serine palmitoyltransferase, mTORC2 has been shown to control the expression of Sptlc1, a subunit of serine palmitoyltransferase, at the level of transcription, leading to changes in complex sphingolipid levels.^73,74^ Second, C17iso-glucosylceramide, one of the complex sphingolipids in *Caenorhabditis elegans*, was shown to promote TORC1 activation.^75,76^ However, opposite results suggesting that glucosylceramide levels appear to be negatively correlated with mTORC1 activity have likewise been reported in Drosophila and mammalian cells as well as *Caenorhabditis elegans*.^77–79^ In addition, roles for sphingolipids in TORC1 activation are consistent with studies showing that a ceramide analog PDMP, which is a well-known glucosylceramide synthesis inhibitor, inhibits mTORC1 activity and affects subcellular localization of mTOR and mammalian Ego1 ortholog LAMTOR1/p18 in mammalian cells.^80,81^ Besides sphingolipids, sterols also have been reported to lie upstream of mTORC1.^82,83^ Third, LAMTOR1/p18, which contains putative myristoylation and palmitoylation domains, was shown to be localized to lipid domains (detergent-resistant membranes) of late endosomes^84^ suggesting a possible conserved link between vacuolar/lysosomal Lo domains and EGO complex-mediated TORC1 activation.

In addition to Lo domain components, other factors may link TORC2 and TORC1 in mammalian cells. Phosphatidic acid (PA) binds to the FRB domain of mTOR.^85–87^ Even in the absence of amino acids, PA drives the translocation of mTORC1 to the lysosome.^88^ Given that an inactive mTORC2 by Rictor (homologue of yeast Avo3) silencing decreases the intracellular PA levels,^89^ it is possible that mTORC2 positively regulates mTORC1 via PA metabolism. Alternatively, since mTORC2 phosphorylates Akt, which functions upstream of mTORC1,^90^ mTORC2 may be linked to mTORC1 by signaling through protein post-translational modifications and interactions. This interplay between mTORC1 and mTORC2 via Akt phosphorylation has been previously proposed in the INS-1 b-cell line.^91^ Although the role of lipids in integrating two functionally different TORCs remains to be resolved in mammalian cells, this and other studies suggest that the integration of TORC1 and TORC2 is an evolutionarily conserved mechanism that coordinates with lipid metabolism to respond to various environmental changes.

Several observations illustrate the interconnections between organelle structure and function and lipid changes. Genes involved in the maintenance of vacuolar structure and function exhibited synthetic phenotypes with *AUR1* (Figure S3), and the inhibition of IPC synthesis induced vacuolar fragmentation (Figure 1B, 1C, and 1D). The mechanism by which vacuolar fragmentation is caused by reduced levels of sphingolipids is unclear, but considering that ergosterol is required for the Sec18/ATP-dependent step of homotypic vacuole fusion,^92^ it is possible that IPC- and ergosterol-rich Lo domains serve as scaffolds for vacuole membrane association of specific proteins, such as the HOPS tethering complex, which are essential for homotypic vacuole fusion. Consistent with this possibility, a yeast strain lacking C26 very-long chain fatty acid-containing sphingolipids is unable to enrich Rab GTPase Ypt7/Rab7 in the vertex region of vacuoles where it functions with the HOPS effector complex during membrane fusion.^23^Interestingly, SNAREs have also been shown to be concentrated in sterol-rich clusters.^93^ On the other hand, lowered sphingolipid levels did not affect vacuole acidification, ESCRT-dependent protein sorting and endosome-to-vacuole vesicular transport, suggesting that IPC-rich Lo domains are not needed for these processes, while the requirement of sterols for optimal V-ATPase function and post-internalization step of endocytosis is well known.^94,95^ Therefore, some specific functions of sterols may be independent of Lo domains.

Conversely, the loss of proteins required for vacuolar acidification dramatically reduced the levels of complex sphingolipids, such as IPC-C. This is consistent with a previous report showing reduced steady-state IPC levels in the *vma2*Δmutant, which has a chronic loss of V-ATPase activity.^96^ Moreover, Tani and Toume reported that the *vma2*Δ mutant caused a decrease in the hydroxylation level of sphingoid bases in the ceramide moiety. As deletion of *SUR2*, which is responsible for the hydroxylation of sphingoid bases, reduces *de novo* IPC synthesis,^18^ a possible explanation for the lowered IPC levels in mutants defective in vacuole acidification may be reduced sphingoid base hydroxylation.

Homotypic vacuole fusion in budding yeast requires the acidification function of vacuolar H+-ATPase (V-ATPase) or the physical presence of V_0_ subunits.^97,98^ One of our surprising findings was that, like vacuole acidification mutants, mutants defective in vacuole fusion also reduced *de novo* IPC synthesis. As vacuolar acidification in vacuole fusion-deficient mutants such as the *aur1-1* mutant (Figure S6A and S6B) and HOPS mutants^99,100^ is maintained normally, the decrease in sphingolipid levels caused by impaired vacuole fusion is not due to the dysfunction of V-ATPase. The mechanism underlying the decrease in sphingolipid levels remains to be elucidated; however, alterations in sphingolipid metabolism due to changes in vacuolar homeostasis may affect the formation of the Lo domain of the vacuolar membrane, resulting in modulation of TORC1 activity. This model is consistent with previous reports that TORC1-related signals were noticeably attenuated by mutations in genes involved in vacuolar acidification and fusion.^68–71,101^

## RESOURCE AVAILABILITY

### Lead contact

Further information and requests for resources should be directed to and will be fulfilled by the Lead Contact, Kouichi Funato.

### Materials availability

The plasmids and yeast strains generated in this study have not yet been deposited in any repository. These materials will be available from the lead contact with a completed material transfer agreement upon request.

### Data and code availability

DNA microarray data for wild-type and AbA-treated wild-type cells were deposited under the GEO Accession Number GSE66973 in NCBI. All other study data are included in the article and the supporting information. This study does not report the original code. Any additional information required to reanalyze the data reported in this study is available from the lead contact upon request.

## ACKNOWLEDGMENTS

We thank H. Riezman for yeast *lcb1-100*, *GhLag1* mutant strains and GFP-Cps1 plasmid, M. Tabuchi for *ypk1ts ypk2*Δ, *rho1*, *slm1-1 slm2*Δ, *pkh1 pkh2*Δ, *stt4-4*, *mss4-102*, *pkc1-2* mutant strains, T. Powers for *avo3-30* mutant strain and *YPK1* plasmids, R. Loewith for pRS416-SCH9-5HA (pJU676) (pRS416) plasmid and T. Noda for GFP-Atg8 (pRS316) plasmid. This work was funded by Grants-in-Aid for Scientific Research from the Japan Society for the Promotion of Science, Japan [JP16K07693 to K.F.; 25K18213 to A.I], by KOSÉ Kobayashi Foundation and Shiraishi Foundation of Science Development to M.M., and by DBT/Welcome Trust India Alliance Fellowship (IA/I/21/1/505624) to S.K.

## AUTHOR CONTRIBUTIONS

HD, KE, KN, RH, TO, SY, AH, KN, YY, AO, YS, TS, KH, KS, PS, AI, KM, MM, and NI constructed the strains and plasmids, performed biochemical studies and fluorescent microscopy experiments, and analyzed the data.

MK performed microarray experiments with the help of H.I.

LM performed GP experiments, and PA and AST performed AFM and FLIM analyses with the help of SK.

KF designed the experiments and wrote the manuscript, with input from all the other authors.

## STAR⍰METHODS

Detailed methods are provided in the online version of this paper and include the following.

## KEY RESOURCES TABLE

In a separate Excel file:

## EXPERIMENTAL MODELS AND SUBJECT DETAILS

### Yeast strains

All strains of *S. cerevisiae* used in this study are listed in the Supplemental Table S1. *AUR1* and *aur1* temperature-sensitive mutant cells were created by introducing pRS415-*AUR1* and pRS415-*aur1* plasmids, respectively, into *aur1* deletion mutant cells transformed with pRS416-GAL1-*AUR1*-3HA, followed by a plasmid shuffling procedure based on 5-fluoroorotic acid (5-FOA) counter-selection. Double mutants were constructed by crossing haploid yeast strains containing single-gene mutations in the same backgrounds, sporulation, and subsequent dissection of the spores. The spore genotypes were verified using colony PCR.

## METHOD DETAILS

### Plasmids

pRS415 (CEN, *LEU2*) plasmids with temperature-sensitive (TS) mutant alleles of the *AUR1* gene were isolated from a random library of mutants generated by error-prone PCR. The mutant library was introduced into *aur1* deletion mutant cells transformed with pRS416-GAL1-*AUR1*-3HA (CEN, *URA3*), and transformants were plated on 5-FOA at 25 °C to shuffle out pRS416-GAL1-*AUR1*-3HA. Colony replica plating was performed onto two YPD plates, which were incubated at 25°C and 37°C for 4 days, and mutants that were not able to grow at 37°C were isolated. From the mutant cells, pRS415 plasmids harboring the TS mutant alleles were recovered, temperature sensitivity was confirmed by plasmid shuffling, and the sequences of the mutant alleles were determined. To construct plasmids overexpressing Gtr1, Gtr2, or Vph1, DNA fragments containing the open reading frame were amplified by PCR and cloned into pRS426 (2μ, *URA3*) with a GDP promoter. The plasmid for the expression of Gtr1-GFP was constructed as follows. The BamHI-BglII fragment containing the GFP(S56T) coding sequence from pFA6a-GFP(S65T)-kanMX6 was inserted into the BamHI site of pRS416 (CEN, *URA3*) to obtain pRS416-GFP. Subsequently, the SalI-EcoRI fragment containing own promoter and open reading frame (without stop codon) of Gtr1 was amplified by PCR and cloned into pRS416-GFP to obtain pRS416-Gtr1-GFP. The fragment containing the promoter and Gtr1-GFP was inserted into enzyme-digested pRS415 (CEN, *LEU2*) to obtain pRS415-Gtr1-GFP. To construct the plasmid expressing Ego1-GFP, the SacI-SalI fragment containing own promoter and open reading frame (without stop codon) of Ego1 was amplified by PCR and inserted into the SalI-SacI sites of pGREG600 (CEN, *URA3*).^102^ To contrast the plasmid overexpressing Vph1-GFP (pRS426-Vph1-GFP), the XhoI fragment containing the GFP coding sequence from pGREG600 was inserted into the XhoI site of pRS426 with GDP promoter to obtain pRS426-GFP, and subsequently, the HindIII-SarI fragment containing the open reading frame (without stop codon) of Vph1 was cloned into pRS426-GFP.

### Culture conditions

Strains were grown either in rich YP medium (1% yeast extract, 2% peptone) supplemented with 0.2% adenine and containing 2% glucose (YPD) as a carbon source, in synthetic complete (SC) containing a complete minimal nutrient mix^103^ (0.15% yeast nitrogen base, 0.5% ammonium sulfate, 2% glucose, all amino acids), or synthetic defined (SD) (0.15% yeast nitrogen base, 0.5% ammonium sulfate, 2% glucose) medium supplemented with the appropriate amino acids and bases as nutritional requirements or with 0.1% 5-FOA. For nitrogen starvation experiments, cells were harvested by centrifugation and grown in an SD-based medium lacking a minimal nutrient mix (SD-N). The sensitivity of yeast strains to drugs, pH, and high temperature was determined by spotting fivefold serial dilutions of cells onto YPD or SD plates and incubating for 4-5 days. Heat stress resistance and chronological life span semi-quantitative assays were performed as previously described.^45,104^

### Fluorescence microscopy

Cells expressing GFP-Atg8, Gtr1-GFP, Eog1-GFP, Vph1-mCherry, or GFP-Cps1 were imaged using differential interference contrast (DIC) and fluorescence microscopy. The ratio of the Gtr1-GFP intensity on the vacuole membrane versus the cytoplasmic signal was determined from a line plot analysis using ImageJ software and calculated by dividing the peak value of the signal intensity (corresponding to the vacuole) by the signal intensity at a point two-thirds of the distance from the vacuole membrane to the plasma membrane (corresponding to the cytoplasm). For vacuole visualization, cells were stained with 20 μM FM4-64 (N-(3-Triethylammoniumpropyl)-4-(6-(4-(diethylamino) phenyl) hexatrienyl) pyridinium) for 15 min at 25°C, washed twice with YPD medium, and incubated for 2h at 25°C or 37°C. Vacuolar morphology was quantified by counting the number of cells containing multilobed vacuoles, as described previously.^105^ To monitor the endocytic pathway, cells were incubated with 20 μM FM4-64 for 30 min at 1-3°C. After washing with YPD, the cells were incubated for the indicated times (5, 20, 40, and 80 min) at 25°C and observed under a fluorescence microscope. For quinacrine staining to visualize vacuolar acidification, cells were grown overnight in YPD buffered with 50 mM NaH_2_PO_4_ at pH 7.6, stained with 200 μM quinacrine for 5 min, washed with SD buffered with 50 mM NaH_2_PO_4_ at pH 7.6, and imaged by fluorescence microscopy. To image the microdomains of vacuoles, cells expressing Ego1-GFP, a marker for liquid-ordered (Lo) domain were incubated in glucose-free SD media (glucose -) for 24h at 25°C and observed under a fluorescence microscope as described.^15^

### DNA microarray analysis

Microarray analysis was performed using the Gene Chip Yeast Genome 2.0 Array (Affymetrix) as described.^106^ Total RNA was extracted from the yeast cells with or without aureobasidin A (AbA) treatment (0.075 μg/ml, for 4h). mRNA was purified, biotinylated cRNA was prepared, and hybridized on the GeneChip Yeast Genome 2.0 Array. GeneChips were stained using a Stain Kit (Affymetrix). Data were analyzed using the Operating Software (GCOS) v1.4, using the Affymetrix default analysis settings and global scaling as the normalization method. DNA microarray data were obtained for three independent culture experiments from wild-type and AbA-treated wild-type cells and can be retrieved from Gene Expression Omnibus (GEO) under the accession code GSE66973.

### Southern blotting, northern blotting, and western blotting

Telomere length was measured by southern blotting as described previously.^18^ Yeast genomic DNA digested with *Xho1* was separated, transferred to Hybond N+ membranes, probed with a digoxigenin-labeled Y’ probe,^107^ and visualized using an anti-digoxigenin antibody conjugated to alkaline phosphatase and CDP-Star substrate. RNA levels were analyzed by northern blotting as described.^108^ GFP-Atg8 and cleaved free GFP were analyzed by SDS-PAGE followed by western blotting using a GFP-specific antibody. Phosphorylated Rps6 and Cdc28 (loading control) were analyzed by western blotting using a rabbit monoclonal anti-phosphor-S6 ribosomal protein (Ser235/236) antibody and a mouse monoclonal anti-PSTAIR antibody. To analyze Sch9 phosphorylation, protein extracts from cells expressing Sch9-5HA were treated with 2-nitro-5-thiocyanobenzoic acid,^47^ separated by SDS-PAGE, and immunoblotted. Blots were probed with mouse or rat anti-HA monoclonal antibody and detected by chemiluminescence using a peroxidase-conjugated affinity-purified anti-mouse or -rat IgG antibody. The bands were quantified using ImageJ software.

### Sphingolipid labeling

Sphingolipids were labeled with [^3^H]*myo*-inositol or [^3^H]dihydrosphingosine (DHS) as described previously.^109^ Radiolabeled lipids were extracted, and half of the sample was subjected to mild alkaline hydrolysis (NaOH) to deacylate glycerophospholipids. Lipids were analyzed by thin-layer chromatography (TLC) using solvent system I (chloroform-methanol-0.25% KCl, 55:45:10, vol/vol/vol) for complex sphingolipids labeled with [^3^H]*myo*-inositol, and solvent system II

(chloroform-methanol-4.2N-ammonium hydroxide, 9:7:2, vol/vol/vol) and III (chloroform-methanol-acetic acid, 190:9:1, vol/vol/vol) for ceramides labeled with [^3^H]dihydrosphingosine (DHS), as described.^110^ Radiolabeled lipids were visualized and quantified using an FLA-7000 system.

### Amino acid analysis

Amino acids were extracted from whole cells and vacuolar fractions as described previously.^111^ Amino acid concentrations in each fraction were measured using an automatic amino acid analyzer (JEOL JLC-500/V).

### Preparation of large unilamellar vesicles (LUVs) and giant unilamellar vesicles (GUVs)

Lipids were extracted from wild-type and *GhLag1* mutant cells using chloroform-methanol-water (10:10:3, v/v/v). The pooled lipid-containing organic solutions were desalted by partitioning with n-butanol, and the lipids were dried under a stream of nitrogen gas. LUV were prepared using the gentle hydration method as described.^111^ Briefly, 0.5 mg of lipid was used in each experiment, and laurdan (1-(6-(dimethylamino) naphthalen-2-yl) dodecan-1-one) was added to the lipid sample to yield a final probe:lipid ratio of 1:500. The lipid solutions were then dried under nitrogen gas and lyophilized overnight. The dried films were hydrated and sonicated, followed by five freeze-thaw cycles to generate LUVs. For GUV preparation, 0.5 mg of lipid was used in each experiment, and a 1:200 (Laurdan:lipid) ratio was maintained. GUVs were prepared using the electroformation method in a temperature-controlled custom-made chamber using optically transparent and electrically conductive indium tin oxide--coated glass coverslips as described.^112^ A 100 µL lipid mixture was spin-coated onto the coverslips and subsequently dried. The coverslips were placed within the electroformation cell, and the cell was filled with Tris-MgCl_2_ buffer. Lipids were hydrated within the cell, and a low-frequency alternating current field was applied. The cells were gradually cooled to room temperature before imaging the GUV.

### Atomic Force Microscopy

AFM imaging was carried out in intermittent contact (alternating current) mode, and force spectroscopy was performed in contact mode with an MFP-3D atomic force microscope (Asylum Research, Santa Barbara, CA) as described.^112^ For imaging and force spectroscopy, silicon nitride cantilevers with spring constant of 0.03-0.26 N/m were used. For preparation of supported lipid bilayer (SLBs), LUVs as prepared above were passed through 100 nm Nucleopore® Polycarbonate Track-Etch™ Membranes (Whatman GmbH, Dassel, Germany) to obtain small unilamellar vesicles (SUVs) and the SUVs were deposited on Ted Pella mica (9.9 mm in diameter) glued on a 60-mm Petri dish and incubated for 2 h in a water bath at 65°C. After the sample was thoroughly washed to remove any unfused SUVs, images were recorded at various locations on the mica. Force spectroscopy was performed to calculate the resistance experienced by the cantilever to rupture the bilayer. The cantilever velocity was maintained at 0.7-1µm/s and the applied force was 15 nN. Force maps were recorded at 16×16 pixels.

### Laurdan generalized polarization (GP) and lifetime Measurements

The generalized polarization (GP) values of Laurdan in LUVs were measured using a Varian Eclipse fluorescence spectrophotometer with a temperature controller with an accuracy of ±0.1°C, and calculated as described previously.^112^ Fluorescence lifetime imaging and microscopy (FLIM) were performed on a MicroTime 200 (Picoquant, Gmbh, Germany) using Time-tagged Time-resolved (TTTR) methodology. This setup was attached to an inverted microscope (IX71) equipped with a water immersion objective (UPlan SApo NA 1.2, 60X, WD = 0.28 mm). Laurdan in GUVs was excited using a 405 nm pulsed diode laser. Fluorescence was collected using a 460/60 band pass filter (AHF/Chroma, Germany), and the collected fluorescence signal was directed to a single photon avalanche photodiode (SPAD) detector. The images were acquired with 512×512 pixel resolution. The fluorescence decays were analyzed using the inbuilt software of MicroTime 200 and fitted using iterative reconvolution to a biexponential function.

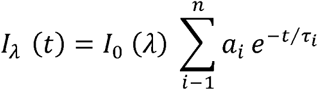

where I(t) is the fluorescence intensity at time t, a_i_ is the pre-exponential factor for the fraction of the fluorescence intensity, τ_i_ is the fluorescence lifetime of the emitting species, and n is the number of exponentials used. The average fluorescence lifetime was calculated using the following equation:

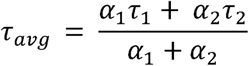

where α1 and α2 are the pre-exponential factors representing the fractional contribution of the decaying component with a lifetime τ_i._

## QUANTIFICATION AND STATISTICAL ANALYSIS

Statistical analysis was performed using Student’s t-test, Tukey-Kramer multiple comparison test, and Fisher’s exact test (ns, not significant; *, p < 0.05; **, p < 0.01; and ***, p < 0.001).

## SUPPLEMENTAL INFORMATION

Supplemental information can be found online at https://////////

## SUPPLEMENTAL FIGURE TITLES AND LEGENDS

**Figure S1.**
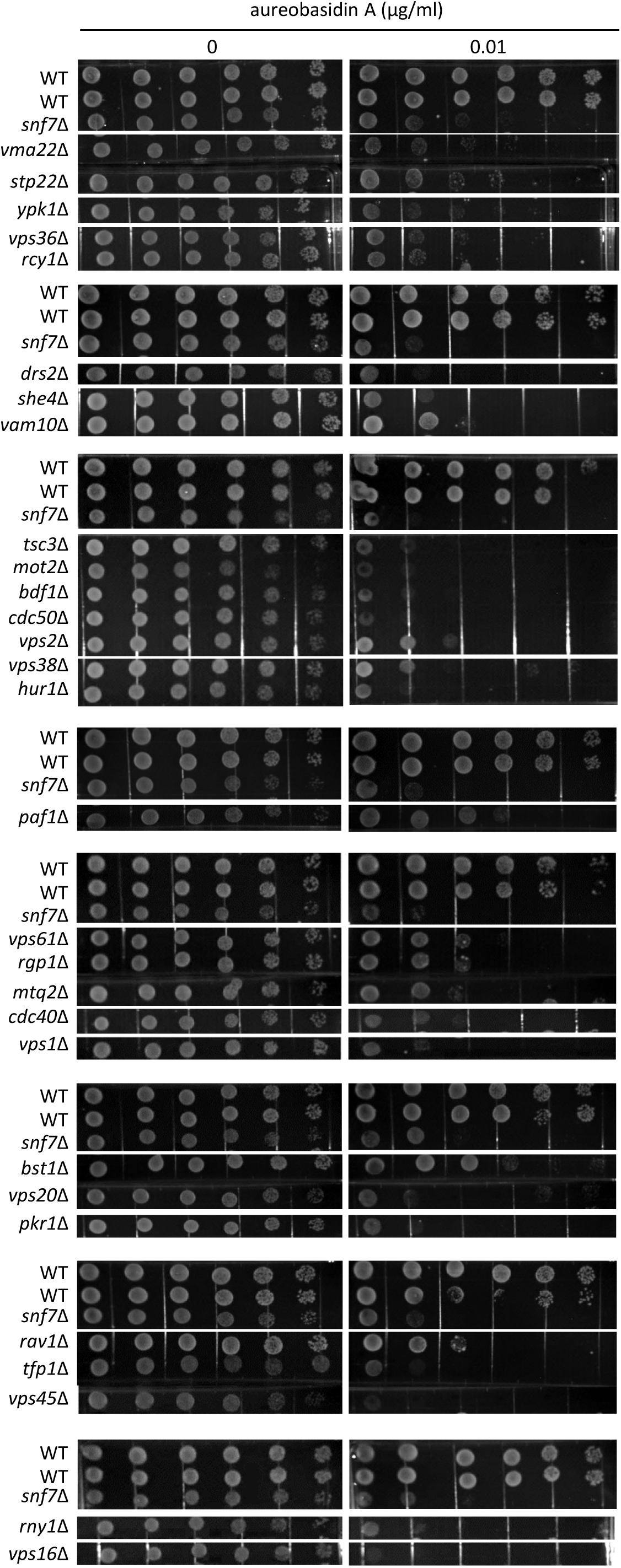
Identification of yeast deletion mutants that confer sensitivity to aureobasidin A (AbA), Related to Fig. 1. Yeast deletion collection haploid Mat-alpha strains were serially diluted fivefold, spotted onto SD plates with or without 0.01 μg/ml AbA, and incubated for 5 days at 25 °C. Of the 4986 yeast deletion strains, 30 strains exhibited exacerbated growth defects caused by AbA.

**Figure S2.**
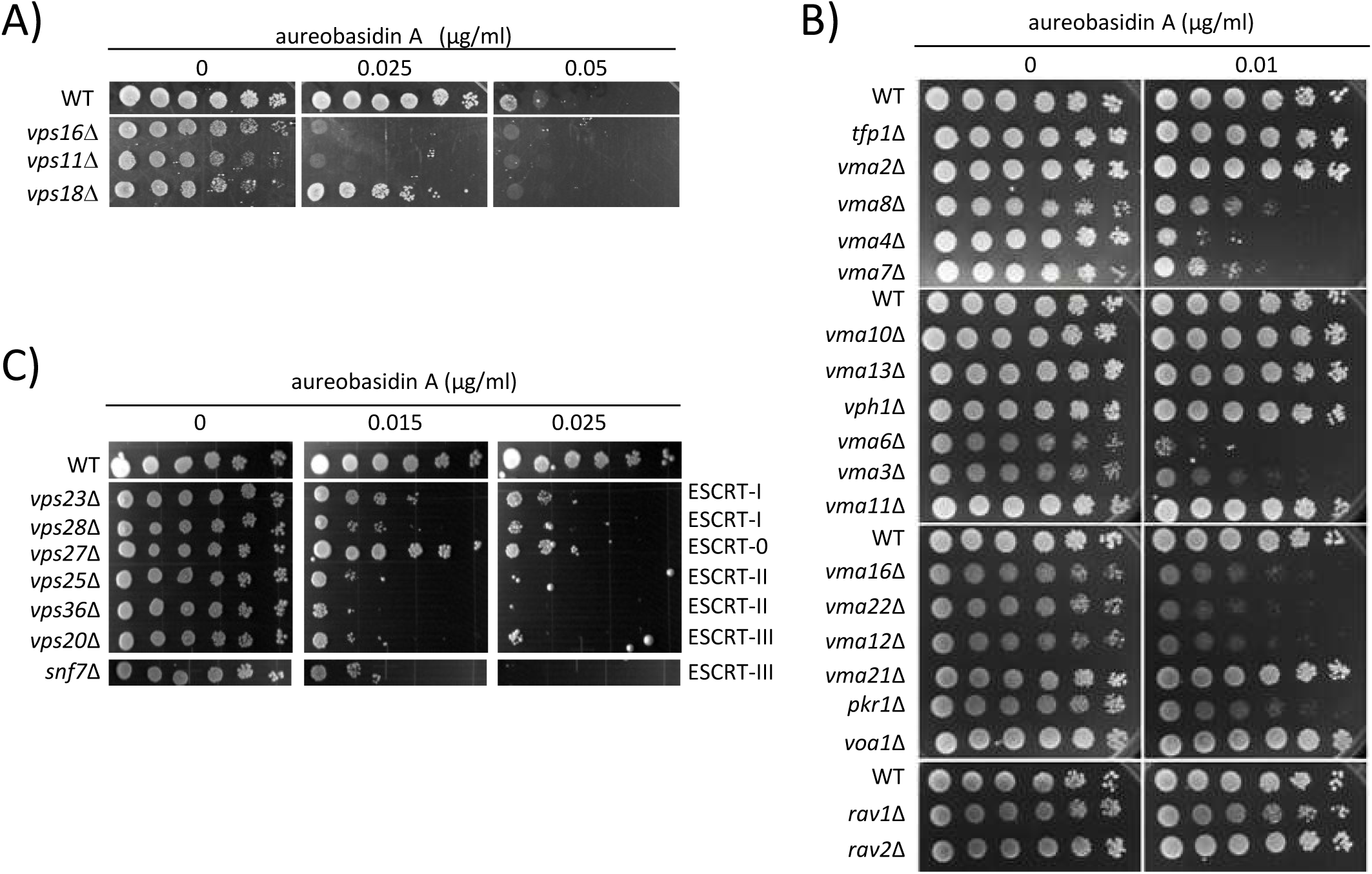
Deletions of genes involved in vacuole fusion, acidification, and ESCRT confer sensitivity to AbA, Related to Fig. 1. Fivefold serial dilutions of deletion mutants for genes involved in vacuole fusion (A), acidification (B), and ESCRT (C) were spotted onto SD plates containing the indicated concentrations of AbA and incubated for 5 days at 25 °C.

**Figure S3.**
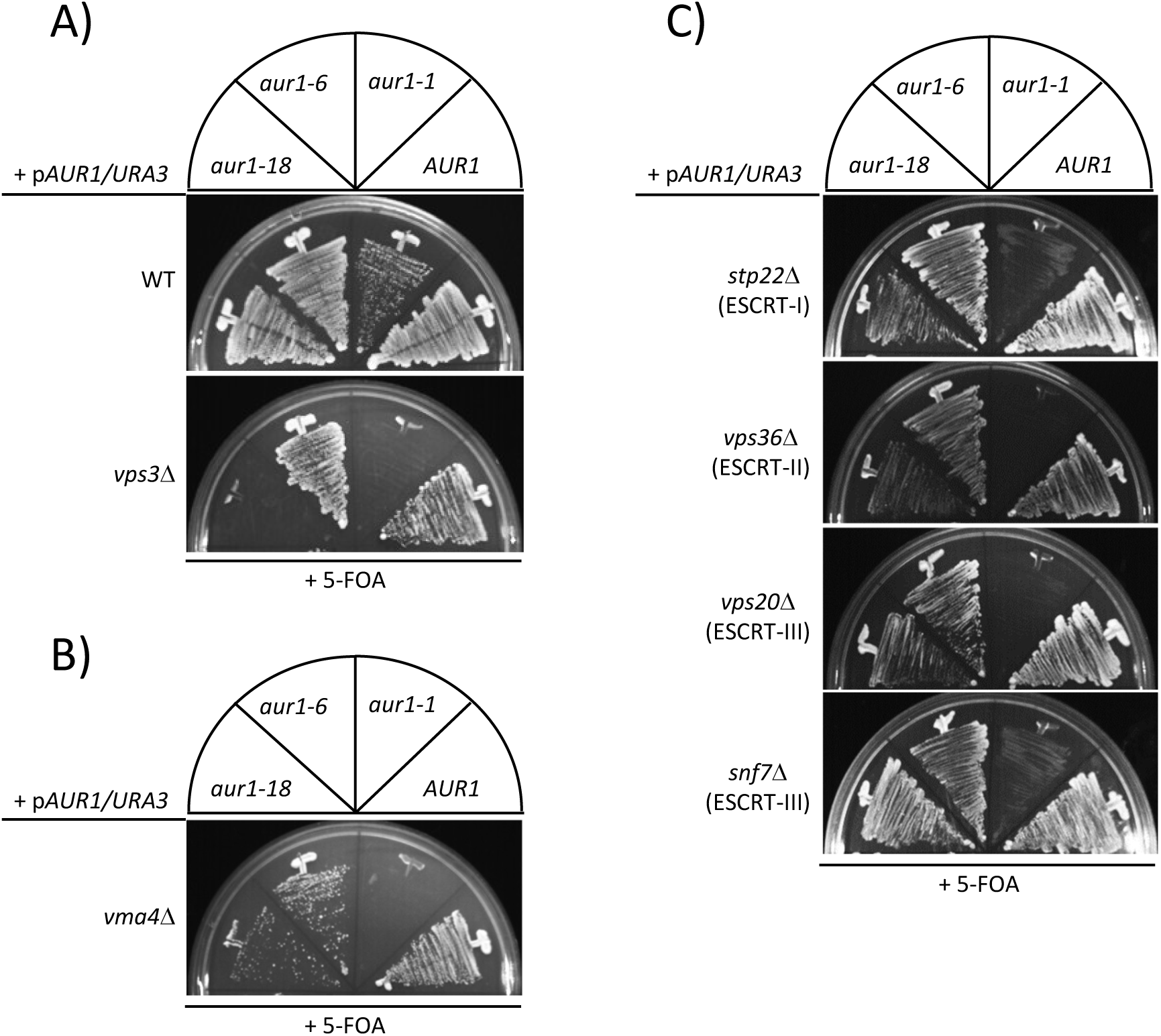
Synthetic lethality occurs between mutations in *AUR1* and genes involved in vacuole fusion, acidification, or ESCRT, Related to Fig. 1. Synthetic lethality between *aur1* mutations and deletion of *VPS3* (A), *VMA4* (B), or ESCRT genes (C) was tested using a plasmid shuffle technique with 5-fluoroacetic acid (5-FOA). *AUR1 xxx*Δ, *aur1-1 xxx*Δ, *aur1-6 xxx*Δ or *aur1-18 xxx*Δ double mutant cells containing pGAL-*AUR1* plasmid with the *URA3* marker (p*AUR1/URA3*) were streaked out on 5-FOA solid medium and allowed to grow at 25°C for 3 days. Failure to grow on 5-FOA plates, which can select cells that have lost the *URA3* plasmid, indicates that the tested double mutant is synthetically lethal.

**Figure S4.**
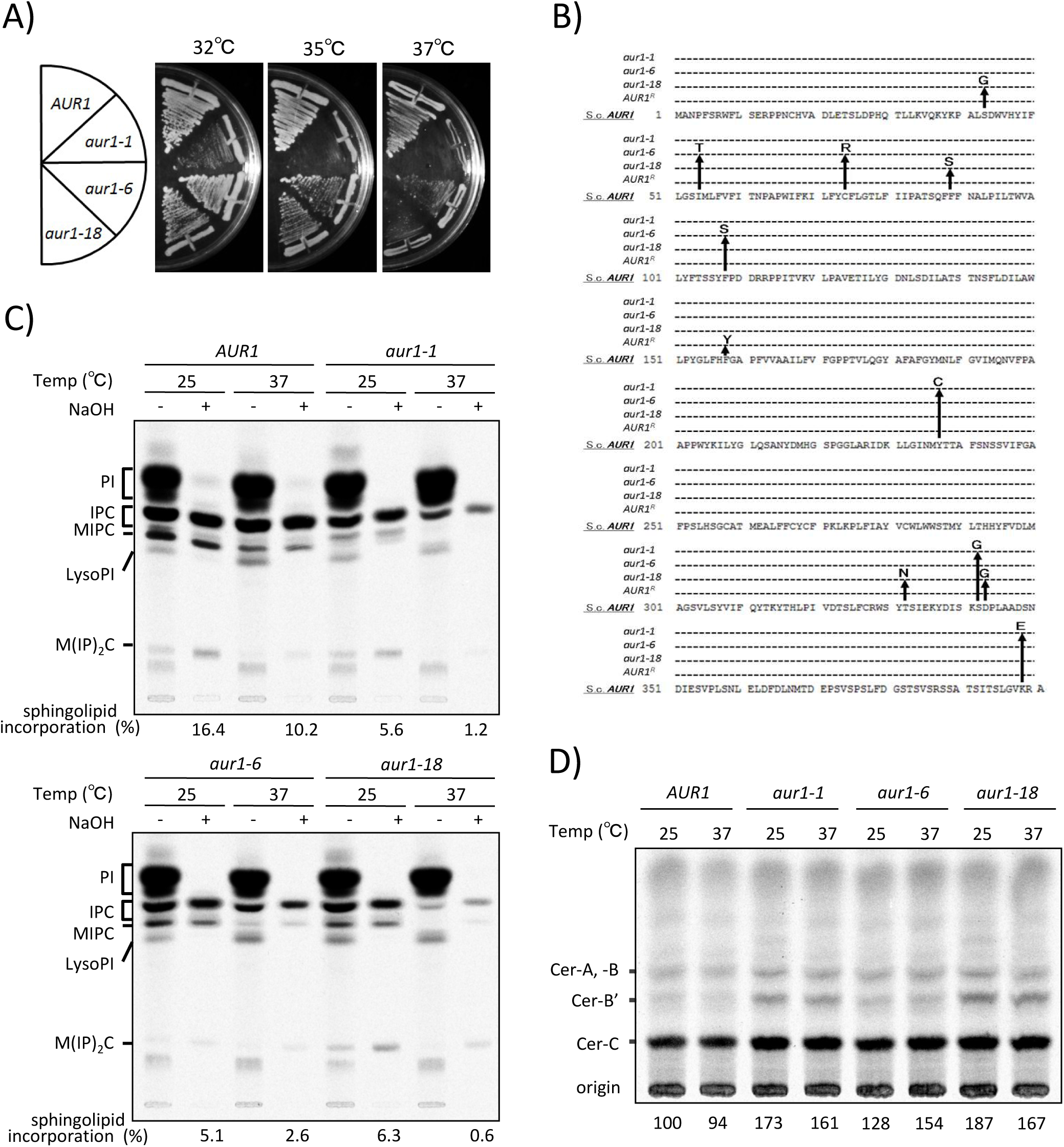
Temperature-sensitive *aur1* mutants have decreased sphingolipid levels and increased accumulation of ceramides, Related to Fig. 1. A) Growth of *AUR1* and *aur1* mutant strains on YPD plates at different temperatures. Temperature-sensitive mutant alleles of *AUR1* were created by random PCR mutagenesis as described previously.^113^ Haploid *AUR1* and *aur1* mutant strains with a chromosomal disruption of the *AUR1* gene and a pRS415 plasmid carrying the *AUR1* or *aur1* mutant alleles *aur1-1*, *aur1-6*, and *aur1-18* were constructed using the plasmid shuffle technique.^114^ *AUR1* and *aur1* mutant cells were grown on YPD plates and incubated at 32, 35, or 37 °C. B) Amino acids substituted in *aur1-1*, *aur1-6*, and *aur1-18* mutated alleles and the AbA-resistant mutant *AUR1R*^115^ are shown. C, D) *AUR1* and *aur1* mutant cells were labeled with [^3^H]*myo*-inositol for 2 h (C) or [^3^H]dihydrosphingosine (DHS) for 1 h (D) at 25 or 37°C. The [^3^H]*myo*-inositol-labeled lipids were treated (+) or not (-) with mild alkaline hydrolysis (NaOH) to differentiate complex sphingolipids from glycerophospholipids and analyzed by thin-layer chromatography (TLC) using solvent system I (C). The incorporation (%) of [^3^H]*myo*-inositol into complex sphingolipids (IPC, MIPC, and M(IP)_2_C) was quantified. The [^3^H] DHS-labeled lipids were treated with NaOH and applied to TLC using solvent system II (D). Fractions containing ceramides on the TLC plate were collected, and the labeled lipids were extracted from the silica and analyzed by second TLC using solvent system III. Total ceramide levels were quantified and presented relative to *AUR1* level at 25°C. The data represent one of two reproducible and independent experiments (C and D).

**Figure S5.**
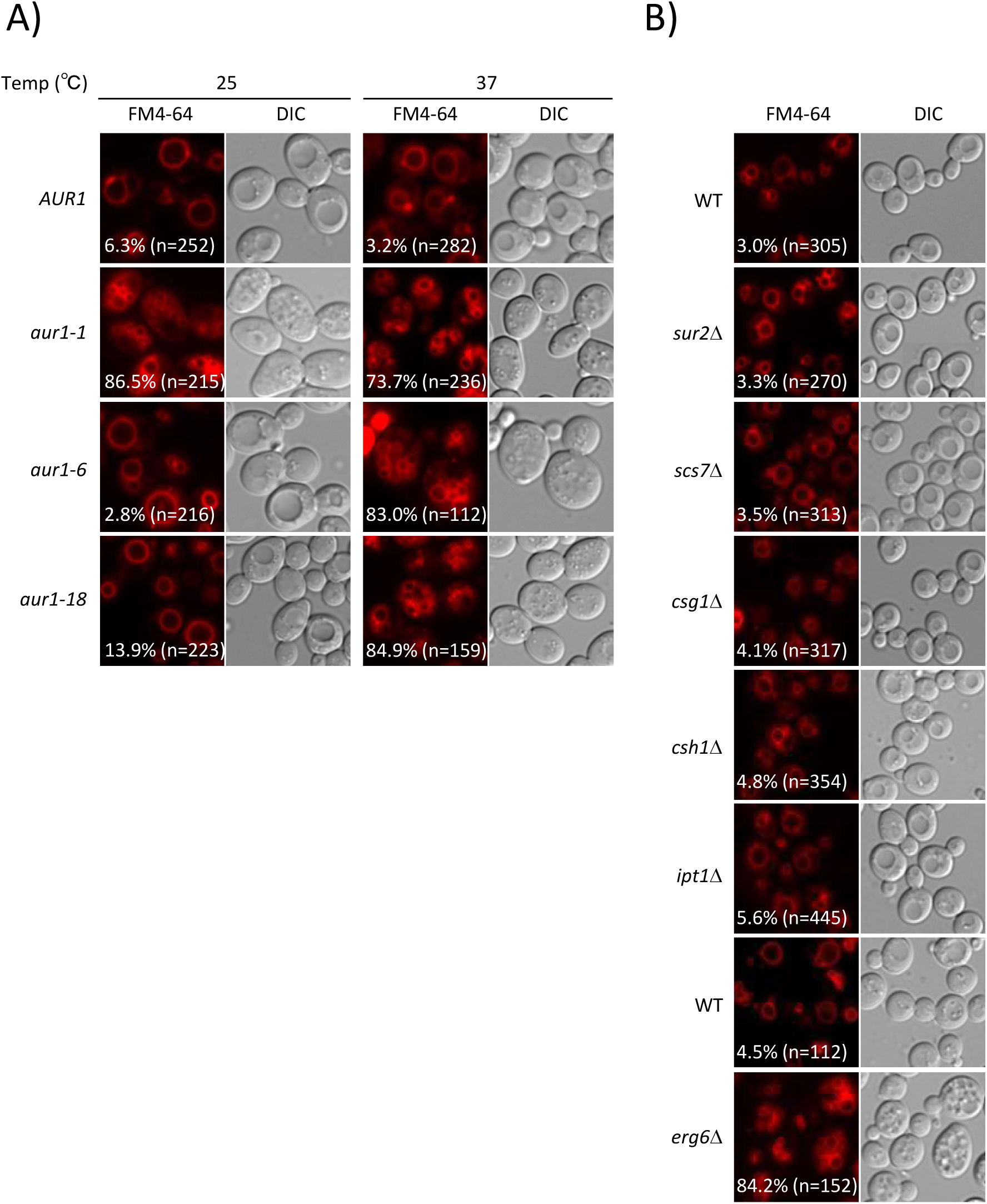
*aur1* mutant cells exhibit highly fragmented vacuolar morphology, Related to Fig. 1. A, B) *AUR1* and *aur1* mutant (A) and wild-type and other mutant (B) cells were grown overnight at 25 °C in YPD medium (A, B), and shifted to 37 °C for 6h (A). Vacuoles were stained with FM4-64 for 2h and imaged using fluorescence microscopy. Vacuole morphology was categorized into round and multi-lobed/fragmented vacuoles as described previously,^105^ and the percentage of cells with multi-lobed/fragmented vacuoles was determined in at least 100 cells (n=112-455). The data represent one of two reproducible independent experiments.

**Figure S6.**
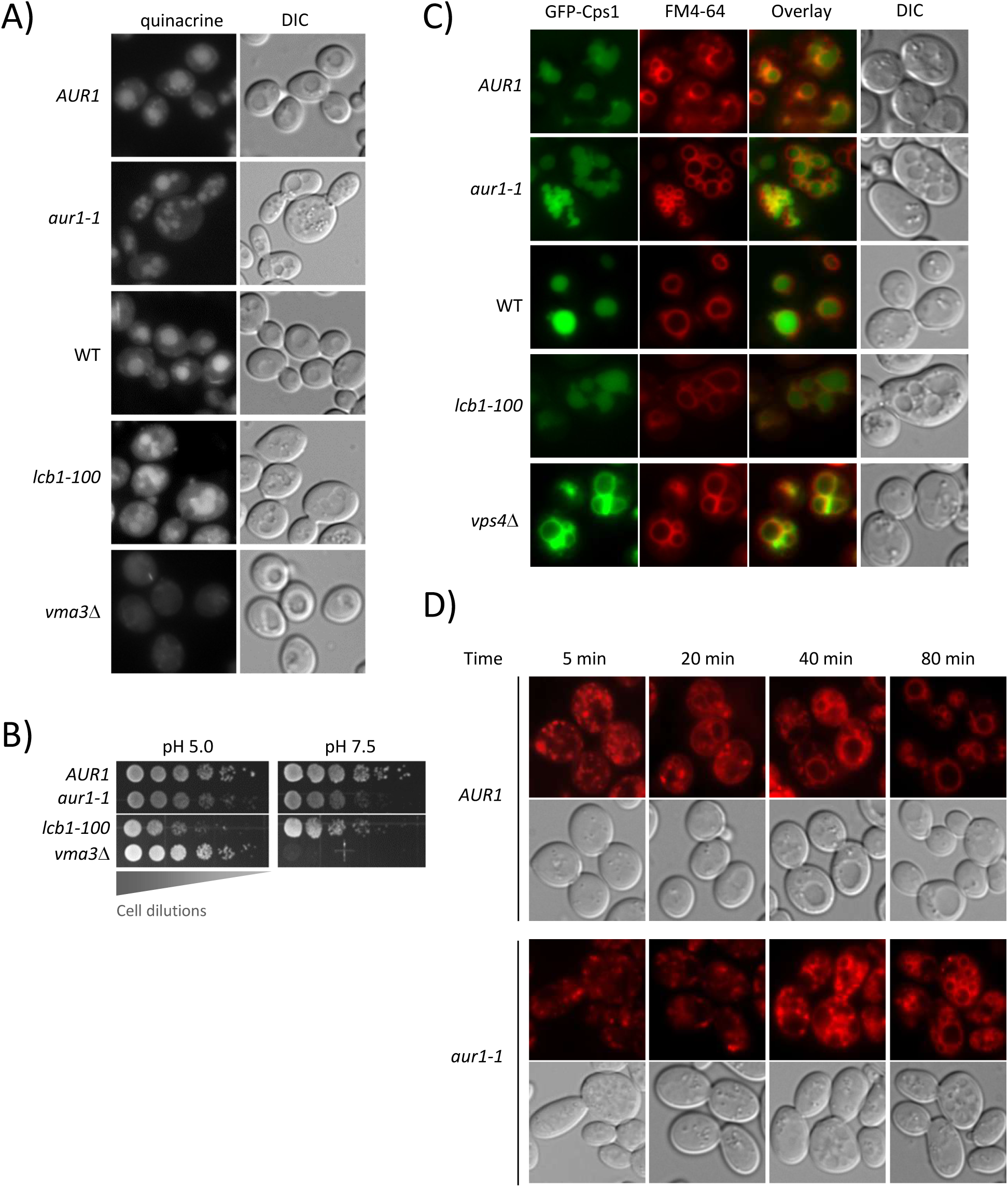
*aur1* mutant cells do not exhibit defects in vacuole acidification, cargo sorting into multivesicular bodies and endocytosis, Related to Fig. 1. A) Cells were grown in YPD buffered at pH 7.6, incubated with quinacrine for 5 min at 25 °C, and then imaged by fluorescence microscopy. B) Fivefold serial dilutions of *AUR1* and mutants were spotted onto pH 5.0 and pH 7.5 YPD plates and incubated for 4 days at 25 °C. C) Cells harboring a plasmid expressing Cps1-GFP were grown in SD medium at 25 °C, stained with FM4-64 for 2h and then imaged by fluorescence microscopy. D) Endocytosis was assessed by tracking FM4-64 uptake as described previously.^94^ Internalization and intracellular transport at 25 °C were monitored at 5, 20, 40, and 80 min.

**Figure S7.**
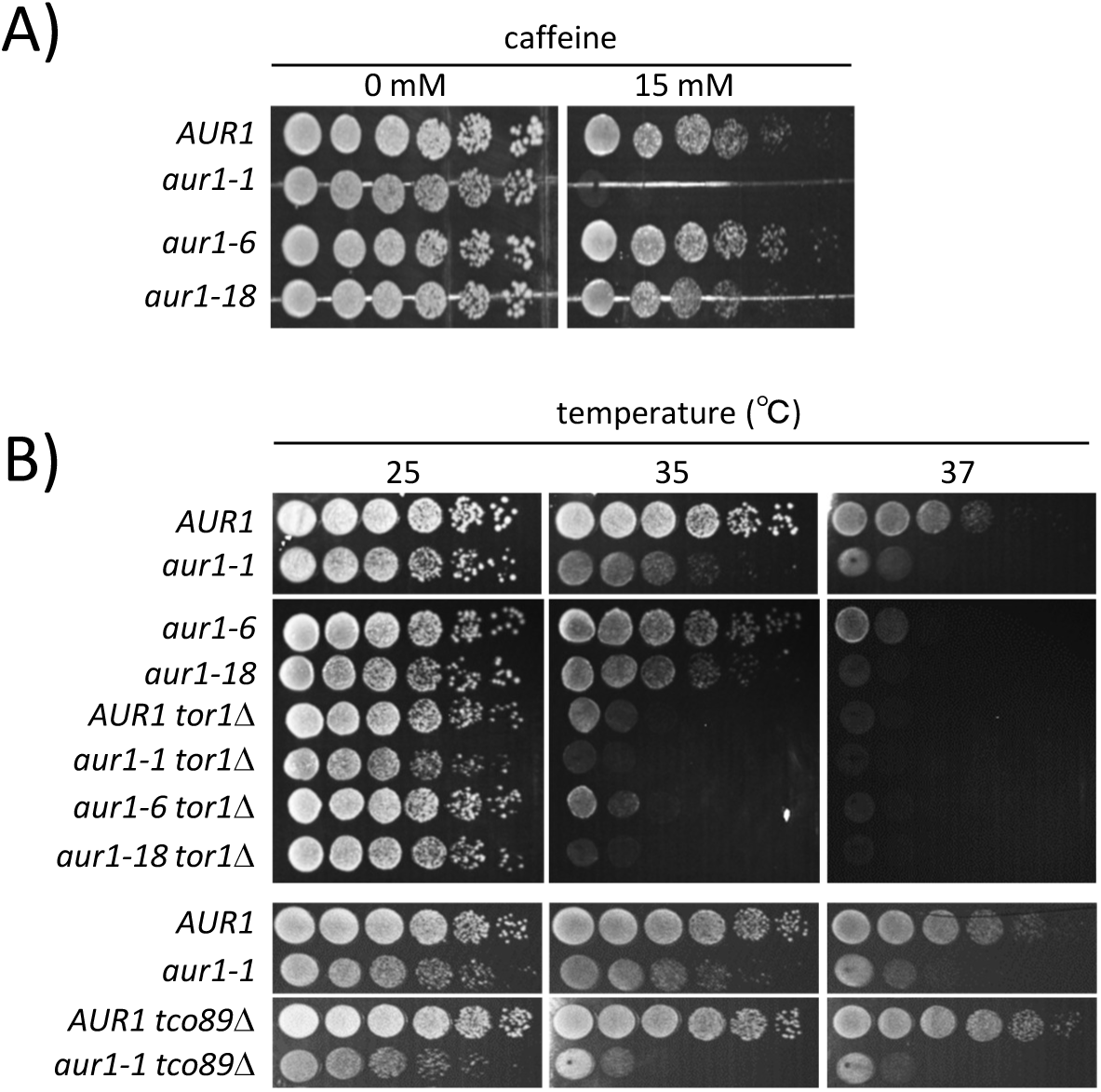
Mutations in *AUR1* confer sensitivity to caffeine, which targets TORC1, and show synthetic growth defects with deletion mutations of TOR1 or TCO89, Related to Fig. 3. A) Hypersensitivity of *aur1-1* mutant cells to caffeine, an inhibitor of TORC1. Cells were serially diluted fivefold, spotted onto YPD plates containing the indicated concentrations of caffeine, and incubated for 4 days at 25 °C. B) Negative genetic interactions between *AUR1* and *TOR1* or *TCO89*. Fivefold serial dilutions of cells were spotted onto YPD plates and incubated for 4 days at the indicated temperatures.

**Figure S8.**
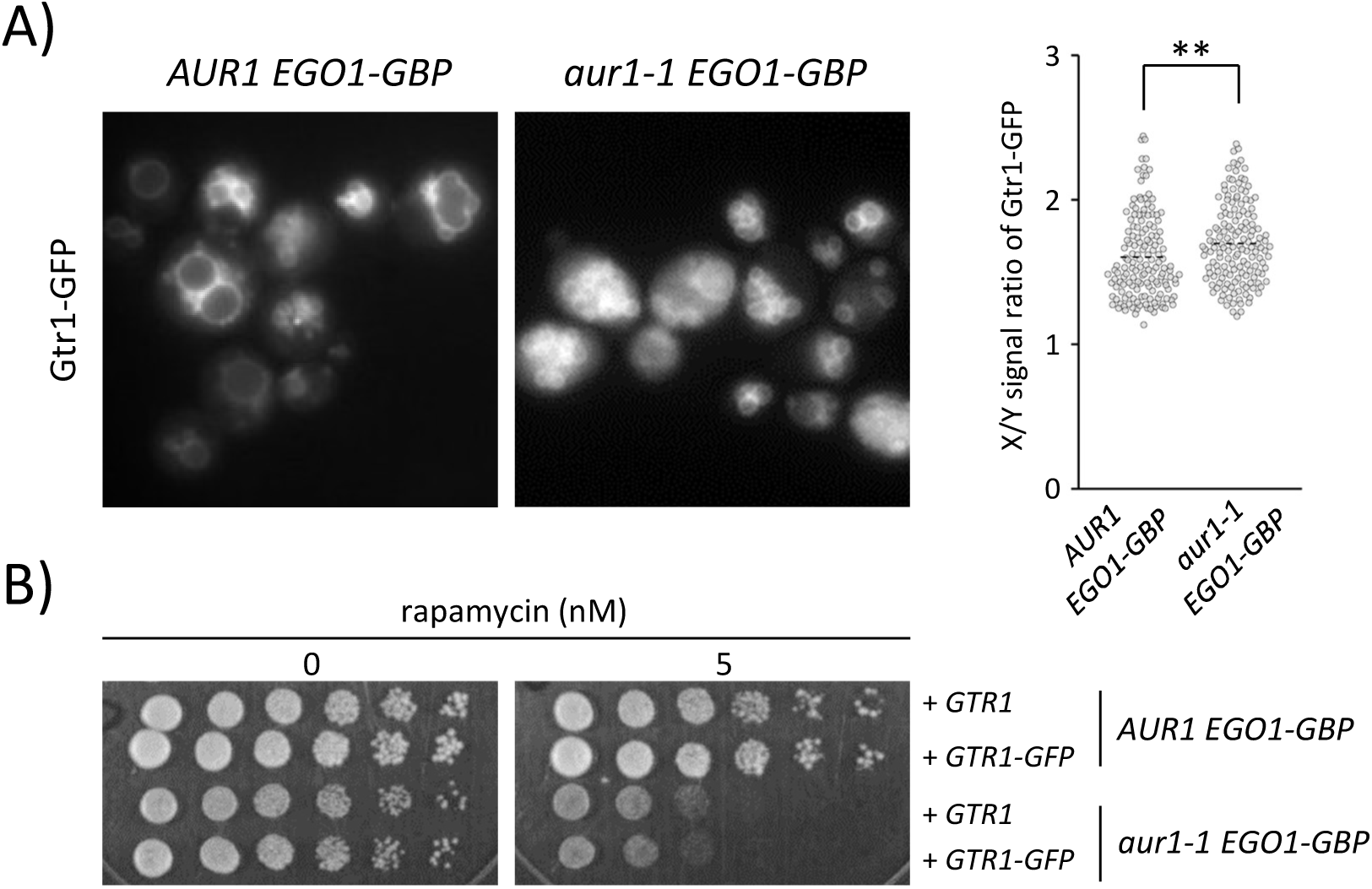
Artificial tethering via interaction with Ego1-GBP rescues the mislocalization of Gtr1-GFP to the cytosol but not rapamycin hypersensitivity in *aur1-1* mutant cells, Related to Fig. 4. A) *AUR1* and *aur1-1* mutant cells expressing Ego1-GBP were transformed with the Gtr1-GFP plasmid and imaged using fluorescence microscopy. The X/Y intensity ratios of Gtr1-GFP were quantified, and the data are represented with beeswarm plots, as described in Fig. 4B. Student’s t test: ***p ≤ 0.001. B) Rapamycin sensitivity. Fivefold serial dilutions of cells were spotted onto YPD plates containing the indicated concentrations of rapamycin and incubated for 4 days at 25 °C.

**Figure S9.**
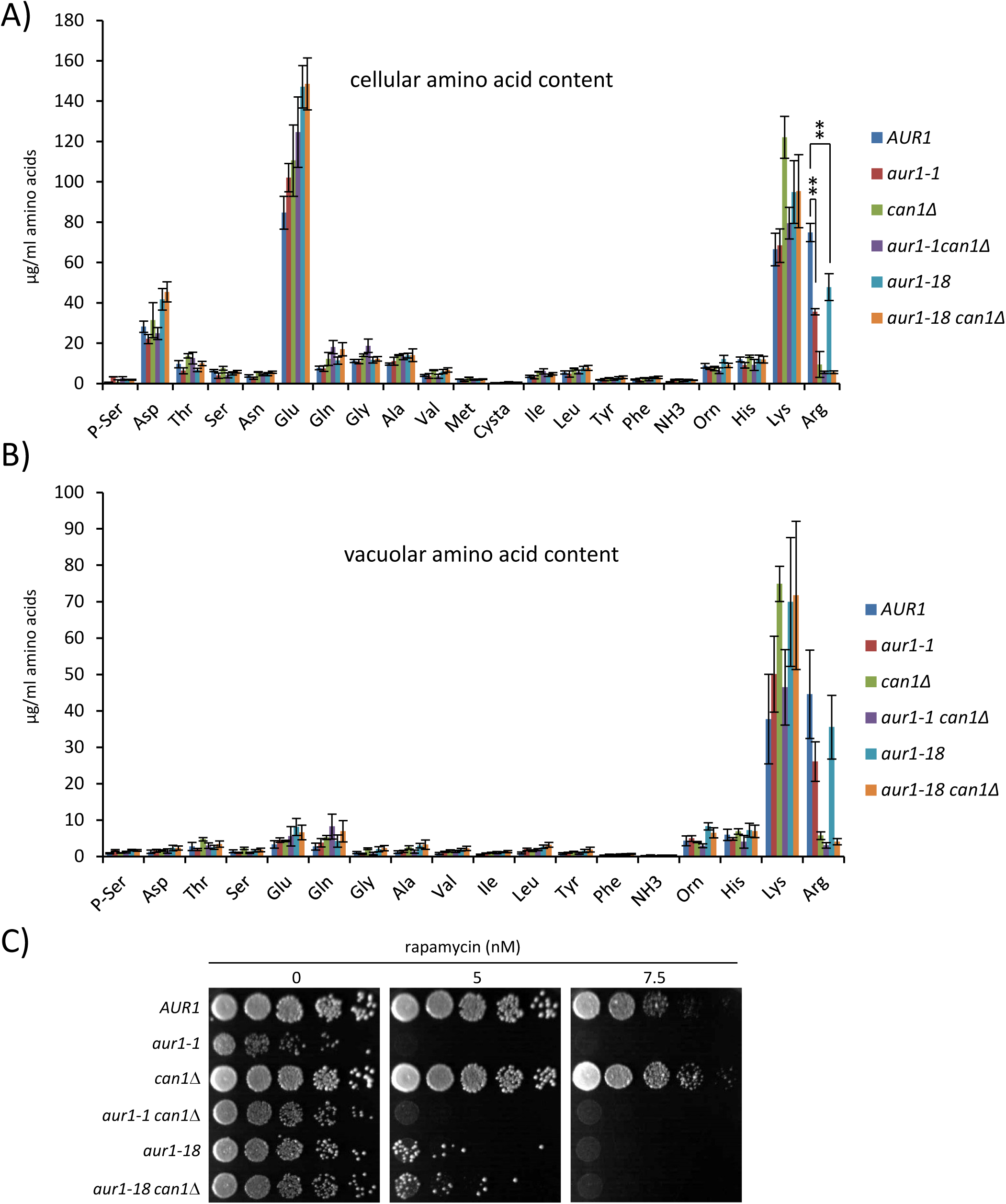
The decrease in arginine levels in *aur1* mutants is not responsible for reduced TORC1 activity, Related to Fig. 4. A, B) *AUR1* and mutants were grown in YPD, and the whole cell (A) and vacuolar (B) amino acid pools were analyzed using an amino acid analyzer. The results are presented as mean ± standard deviation (SD) based on at least three independent experiments. Student’s t test: **p ≤ 0.01. C) Rapamycin sensitivity. Fivefold serial dilutions of cells were spotted onto YPD plates containing the indicated concentrations of rapamycin and incubated for 4 days at 25 °C.

**Figure S10.**
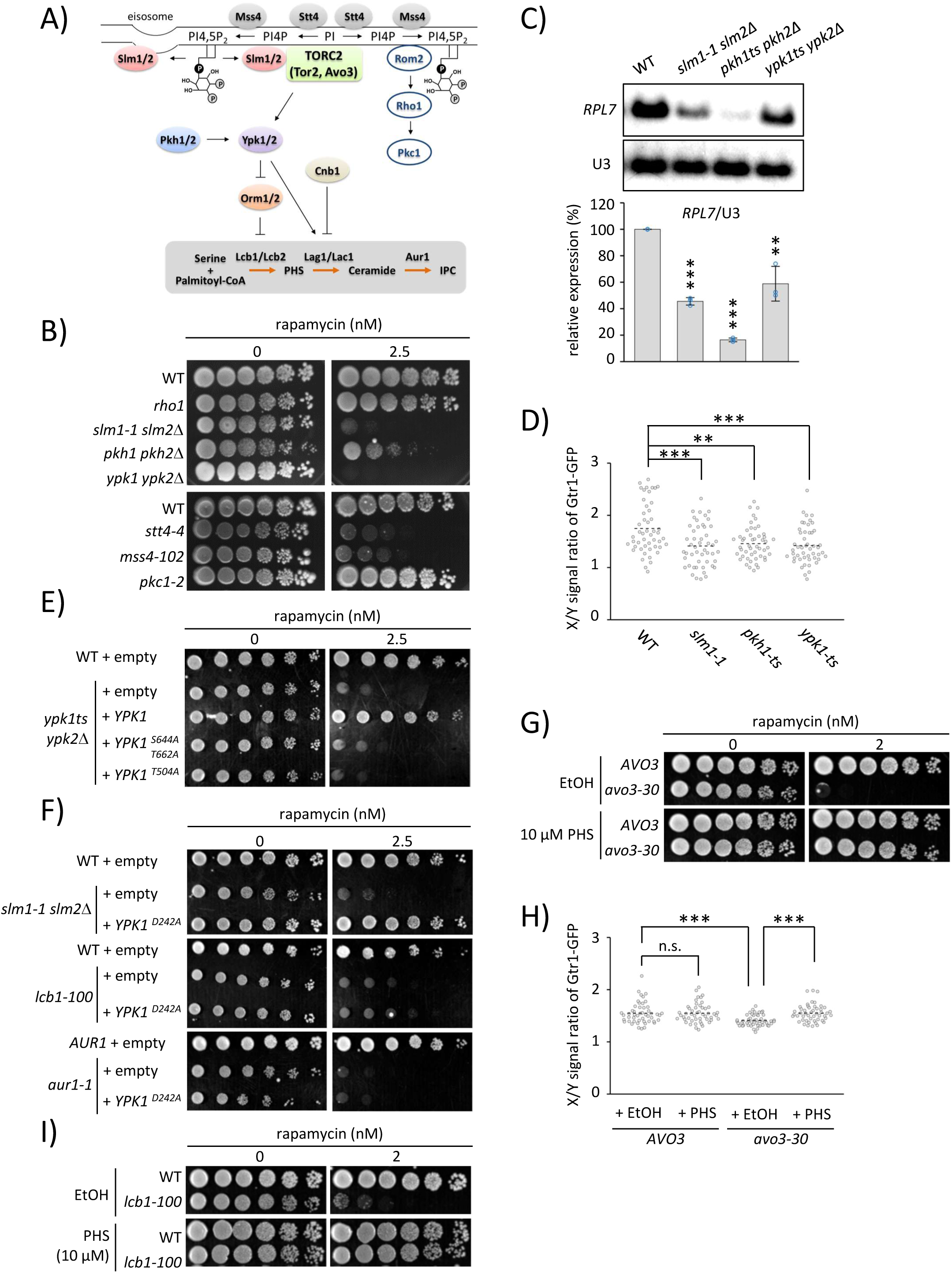
TORC2-Ypk signaling regulates TORC1 activation via sphingolipid metabolism, Related to Fig. 6. A) Schematic representation of the regulation of sphingolipid synthesis by TORC2-Ypk signaling. TORC2 regulates sphingolipid biosynthesis via the direct phosphorylation of Ypk1/2. Ypk1/2 kinases are phosphorylated and activated by Pkh1/2 kinases. Mss4-dependent release of Slm1 from MCC/eisosomes stimulates TORC2 activation, which mediates Ypk1/2 phosphorylation. The TORC2-Ypk1/2 signaling pathway phosphorylates Orm1/2 proteins to derepress sphingolipid synthesis, whereas the regulatory subunit of calcineurin, Cnb1, downregulates ceramide synthase activity by opposing the TORC2-Ypk1/2 signaling pathway. The generation of PI4,5P2 by the sequential actions of Stt4 and Mss4 recruits Rom2 to the plasma membrane, which activates the Rho1-Pkc1 cascade for cell wall integrity (CWI) signaling. B, E, F, G, and I) Rapamycin sensitivity of wild-type, *AVO3,* and mutants was determined by spotting serial dilutions of cells onto YPD plates with or without the indicated concentrations of rapamycin (B). Wild-type and mutant cells transformed with empty plasmids (E, F) or plasmids expressing wild-type Ypk1, Ypk1S664A/T662A, Ypk1T504A (E), and Ypk1D242A (F) were spotted onto SC plates lacking leucine and methionine (SC-LM) (E) and SC-UM plates (F) with or without 2.5 nM rapamycin. *AVO3*, wild-type, and mutant cells were spotted onto YPD plates with 10 μM PHS or ethanol as a vehicle control, with or without 2 nM rapamycin (G, I). Then, cells were grown for 4 days at 25 °C. C) Levels of *RPL7* mRNA and U3 snoRNA (as a loading control) were analyzed by northern blotting, quantified from three independent experiments, normalized to U3, and expressed as the mean ± SD of relative levels. Student’s t test: **p < 0.01; ***P < 0.001. D, H) Cells expressing Gtr1-GFP treated without (D) and with 10 μM PHS or ethanol (H) were observed by fluorescence microscopy, and X/Y intensity ratios were quantified and data are represented with beeswarm plots as described in Fig. 4B. Student’s t-test: **p < 0.01; ***P < 0.001 (D), Tukey-Kramer multiple comparison test ***P < 0.001 (H).

**Figure S11.**
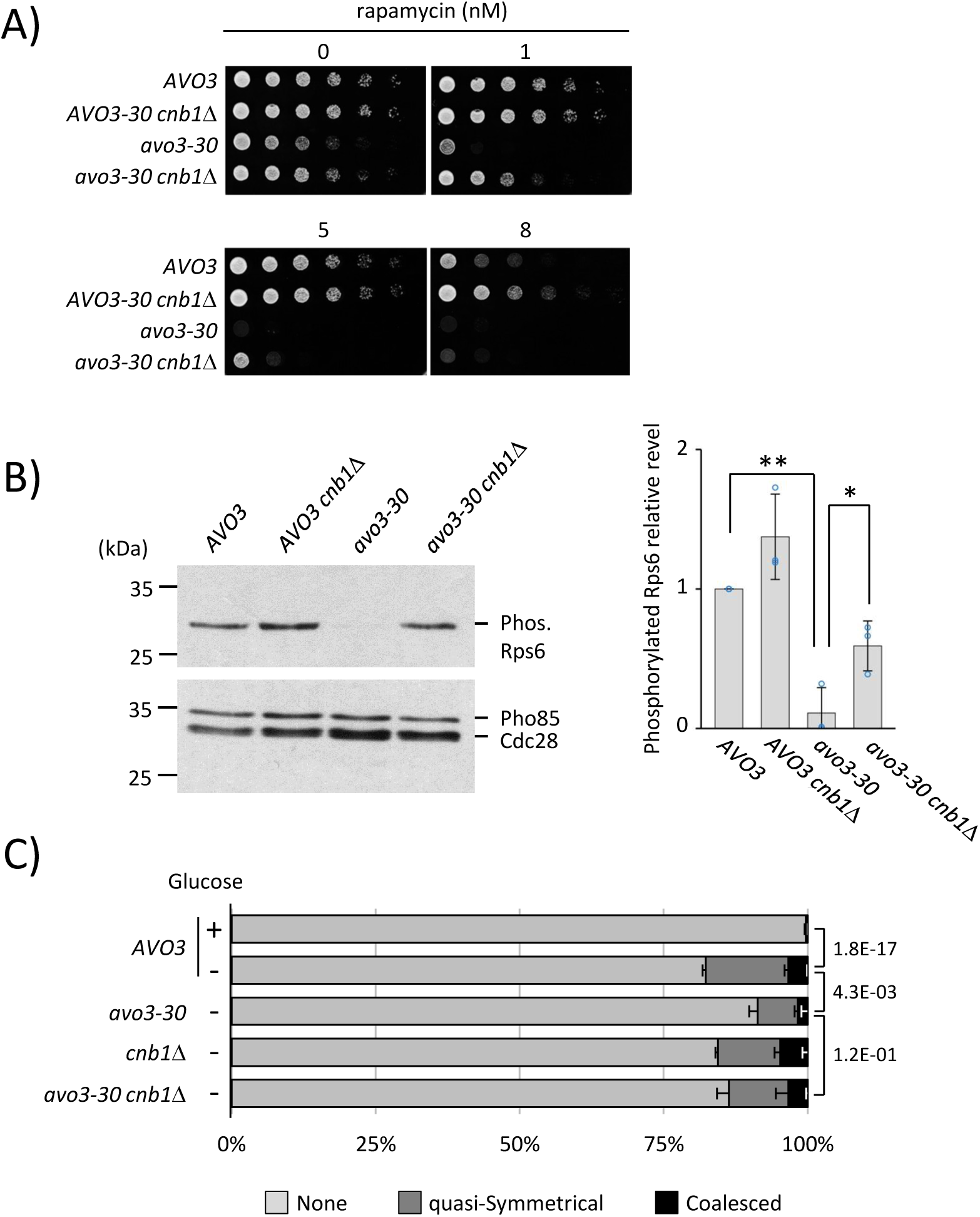
Calcineurin antagonizes the TORC2-Ypk1/2-dependent regulation of TORC1, Related to Fig. 6. A) Rapamycin sensitivity of *AVO3* and mutants was determined by spotting serial dilutions of cells onto YPD plates with or without the indicated concentrations of rapamycin. B) Western blotting and quantification of Rps6 phosphorylation levels. The levels of phosphorylated Rps6 (a proxy for TORC1 activity) and Cdc28 (loading control) were analyzed using western blotting. The ratio of phosphorylated Rps6 and Cdc28 was normalized to that in *AVO3* control cells, and relative levels were expressed as the mean ± SD from three independent experiments. Tukey-Kramer multiple comparison test: **p ≤ 0.01; *p ≤ 0.05. C) *AVO3* and mutant cells expressing Ego1-GFP were incubated in SD or SD lacking glucose at 25°C for 24 h, as described in Fig 4D. The percentage of cells with vacuolar domains was calculated from 100 cells and expressed as the mean ± SD of three independent experiments. Statistical significance was tested using Fisher’s exact test.

**Figure S12.**
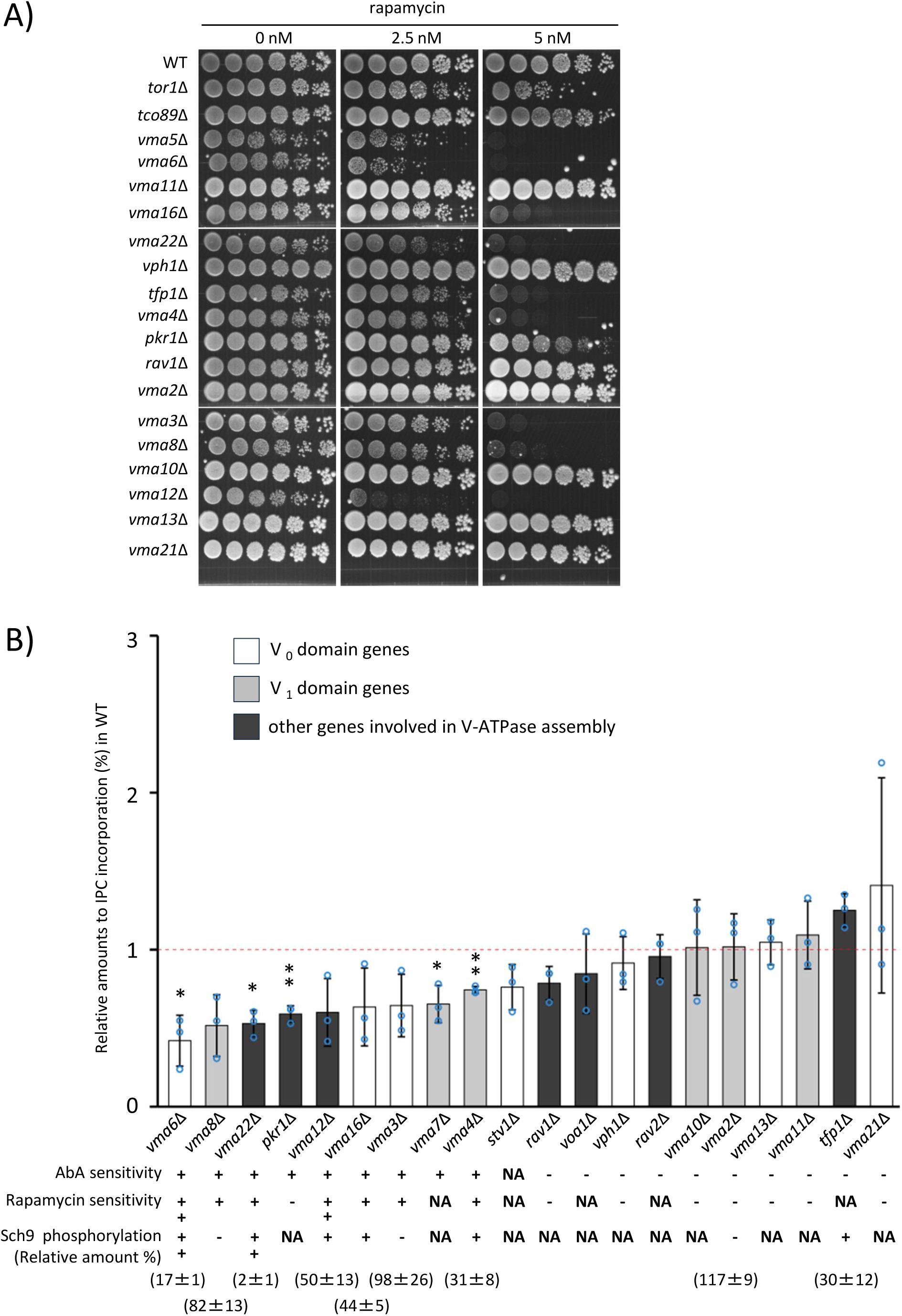
V-ATPase regulates TORC1 activation via IPC synthesis, Related to Fig. 7. A) Rapamycin sensitivity of deletion mutants of V-ATPase genes was assessed on YPD containing the indicated concentrations of rapamycin at 25 °C. B) Incorporation (%) of [^3^H]*myo*-inositol into IPC (for 90 min at 25 °C) and Sch9 phosphorylation were analyzed as described in Fig. 7D and Fig. 3C, respectively. The relative amounts of incorporation (%) into IPC and phosphorylated (Sch9-P)/total Sch9 ratio to wild-type cells were determined from three independent experiments. Student’s t test: *p < 0.05; **P < 0.01. Cells showing a strong reduction in Sch9 phosphorylation and hypersensitivity to drugs (AbA and rapamycin) are indicated with two plus signs (+ +). A plus sign (+) indicates that cells have a mild reduction in the phosphorylation of Sch9 and are sensitive to drugs, and a minus sign (-) indicates that there is no obvious difference between the deletion mutant and wild-type cells. NA; not analyzed. The data on sensitivity to AbA and rapamycin are provided in supplemental Figure S2 (B) and Figure S12 (A), respectively.

**Figure S13.**
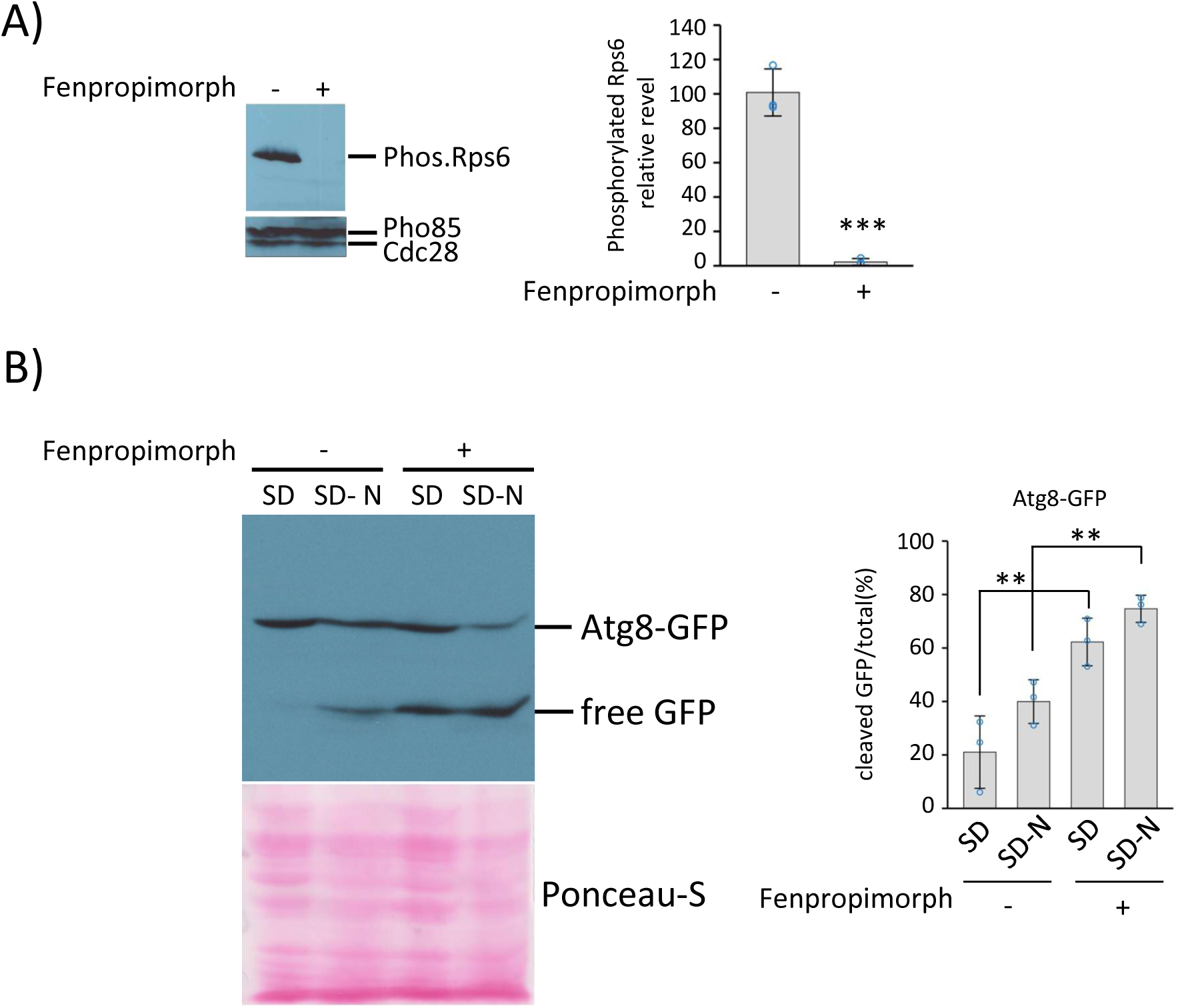
Sterols are required for TORC1 activation, Related to Fig. 3. A) The levels of phosphorylated Rps6 and Cdc28/Pho84 (loading control) were analyzed by western blotting after overnight treatment with 1μM fenpropimorph (left panel). The ratio of phosphorylated Rps6 to Cdc28/Pho84 was normalized to that in untreated control cells, and relative levels are expressed as the mean ± SD from three independent experiments (right panel). Student’s t test: ***p ≤ 0.001. B) The cells expressing GFP-Atg8 were grown exponentially in SD with or without 100μM fenpropimorph, and then starved for nitrogen (120 min; SD-N). Autophagic degradation of GFP-Atg8 was analyzed by western blotting (left panel), and autophagic flux is expressed as the mean ± SD from three independent experiments (right panel). Tukey-Kramer multiple comparison test: **p ≤ 0.01.

## SUPPLEMENTAL TABLE TITLE

**Supplemental Table S1.** Strains of *Saccharomyces cerevisiae* used for this study.

## REFERENCES

1. Cohen, S., Valm, A.M., and Lippincott-Schwartz, J. (2018). Interacting organelles.Curr. Opin. Cell Biol. 53, 84–91. doi: 10.1016/j.ceb.2018.06.003.

2. Cui, L., Li, H., Xi, Y., Hu, Q., Liu, H., Fan, J., Xiang, Y., Zhang, X., Shui, W., and Lai, Y. (2022). Vesicle trafficking and vesicle fusion: mechanisms, biological functions, and their implications for potential disease therapy. Mol. Biomed. 3, 29. doi: 10.1186/s43556-022-00090-3.

3. Farhan, H., and Rabouille, C. (2011). Signalling to and from the secretory pathway. J. Cell Sci. 124, 171–80. doi: 10.1242/jcs.076455.

4. Jain, A., and Zoncu, R. (2022). Organelle transporters and inter-organelle communication as drivers of metabolic regulation and cellular homeostasis. Mol. Metab. 60, 101481. doi: 10.1016/j.molmet.2022.101481.

5. Loewith, R., and Hall, M.N. (2011). Target of rapamycin (TOR) in nutrient signalling and growth control. Genetics 189, 1177–201. doi: 10.1534/genetics.111.133363.

6. Prinz, W.A., Toulmay, A., and Balla, T. (2020). The functional universe of membrane contact sites. Nat. Rev. Mol. Cell Biol. 21, 7–24. doi: 10.1038/s41580-019-0180-9.

7. Teixeira, V., and Costa, V. (2016). Unraveling the role of the Target of Rapamycin signalling in sphingolipid metabolism. Prog. Lipid Res. 61, 109–33. doi: 10.1016/j.plipres.2015.11.001.

8. Levental, I., Levental, K.R., and Heberle, F.A. (2020). Lipid Rafts: Controversies Resolved, Mysteries Remain. Trends Cell Biol. 30, 341–353. doi: 10.1016/j.tcb.2020.01.009.

9. Malinsky, J., Opekarová, M., and Tanner, W. (2010). The lateral compartmentation of the yeast plasma membrane. Yeast 27, 473–8. doi: 10.1002/yea.1772.

10. Murley, A., Yamada, J., Niles, B.J., Toulmay, A., Prinz, W.A., Powers, T., and Nunnari, J. (2017). Sterol transporters at membrane contact sites regulate TORC1 and TORC2 signalling. J. Cell Biol. 216, 2679–2689. doi: 10.1083/jcb.201610032.

11. Spira, F., Mueller, N.S., Beck, G., von Olshausen, P., Beig, J., and Wedlich-Söldner, R. (2012). Patchwork organization of the yeast plasma membrane into numerous coexisting domains. Nat. Cell Biol. 14, 640–8. doi: 10.1038/ncb2487.

12. Toulmay, A., and Prinz, W.A. (2013). Direct imaging reveals stable, micrometer-scale lipid domains that segregate proteins in live cells. J. Cell Biol. 202, 35–44. doi: 10.1083/jcb.201301039.

13. Wang, C.W., Miao, Y.H., and Chang, Y.S. (2014). A sterol-enriched vacuolar microdomain mediates stationary phase lipophagy in budding yeast. J. Cell Biol. 206, 357–66. doi: 10.1083/jcb.201404115.

14. Kim, H., and Budin, I. (2024). Intracellular sphingolipid sorting drives membrane phase separation in the yeast vacuole. J. Biol. Chem. 300, 105496. doi: 10.1016/j.jbc.2023.105496.

15. Sakuragi, K., Schlarmann, P., Ikeda, A., and Funato K. (2023). Vacuole membrane contact sites regulate liquid-ordered domain formation during glucose starvation. FEBS Lett. 597, 1462–1468. doi: 10.1002/1873-3468.14621.

16. Rahman, M.A., Kumar, R., Sanchez, E., and Nazarko, T.Y. (2021). Lipid Droplets and Their Autophagic Turnover via the Raft-Like Vacuolar Microdomains. Int. J. Mol. Sci. 22, 8144. doi: 10.3390/ijms22158144.

17. Numrich, J., Péli-Gulli, M.P., Arlt, H., Sardu, A., Griffith, J., Levine, T., Engelbrecht-Vandré, S., Reggiori, F., De Virgilio, C., and Ungermann, C. (2015). The I-BAR protein Ivy1 is an effector of the Rab7 GTPase Ypt7 involved in vacuole membrane homeostasis. J. Cell Sci. 128, 2278–92. doi: 10.1242/jcs.164905.

18. Ikeda, A., Muneoka, T., Murakami, S., Hirota, A., Yabuki, Y., Karashima, T., Nakazono, K., Tsuruno, M., Pichler, H., Shirahige, K., Kodama, Y., Shimamoto, T., Mizuta, K., and Funato, K. (2015). Sphingolipids regulate telomere clustering by affecting the transcription of genes involved in telomere homeostasis. J. Cell Sci. 128, 2454–67. doi: 10.1242/jcs.164160.

19. Kajiwara, K., Muneoka, T., Watanabe, Y., Karashima, T., Kitagaki, H., and Funato, K. (2012). Perturbation of sphingolipid metabolism induces endoplasmic reticulum stress-mediated mitochondrial apoptosis in budding yeast. Mol. Microbiol. 86, 1246–61. doi: 10.1111/mmi.12056.

20. Murakami, S., Shimamoto, T., Nagano, H., Tsuruno, M., Okuhara, H., Hatanaka, H., Tojo, H., Kodama, Y., and Funato. K. (2015). Producing human ceramide-NS by metabolic engineering using yeast Saccharomyces cerevisiae. Sci. Rep. 5, 16319. doi: 10.1038/srep16319.

21. Zanolari, B., Friant, S., Funato, K., Sütterlin, C., Stevenson, B.J., and Riezman, H. (2000). Sphingoid base synthesis requirement for endocytosis in Saccharomyces cerevisiae. EMBO J. 19, 2824–33. doi: 10.1093/emboj/19.12.2824.

22. Faergeman, N.J., Feddersen, S., Christiansen, J.K., Larsen, M.K., Schneiter, R., Ungermann, C., Mutenda, K., Roepstorff, P., and Knudsen, J. (2004). Acyl-CoA-binding protein, Acb1p, is required for normal vacuole function and ceramide synthesis in Saccharomyces cerevisiae. Biochem. J. 380, 907–18. doi: 10.1042/BJ20031949.

23. Zhang, C., Calderin, J.D., Hurst, L.R., Gokbayrak, Z.D., Hrabak, M.R., Balutowski, A., Rivera-Kohr, D.A., Kazmirchuk, T.D.D., Brett, C.L., and Fratti R.A. (2024). Sphingolipids containing very long-chain fatty acids regulate Ypt7 function during the tethering stage of vacuole fusion. J. Biol. Chem. 300, 107808. doi: 10.1016/j.jbc.2024.107808.

24. Schlarmann, P., Ikeda, A., and Funato, K. (2021). Membrane Contact Sites in Yeast: Control Hubs of Sphingolipid Homeostasis. Membranes (Basel) 11, 971. doi: 10.3390/membranes11120971.

25. Funato, K., and Riezman, H. (2001). Vesicular and nonvesicular transport of ceramide from ER to the Golgi apparatus in yeast. J. Cell Biol. 155, 949–59. doi: 10.1083/jcb.200105033.

26. Sambade, M., Alba, M., Smardon, A.M., West, R.W., and Kane, P.M. (2005). A genomic screen for yeast vacuolar membrane ATPase mutants. Genetics 170, 1539–51. doi: 10.1534/genetics.105.042812.

27. Weisman, L.S., Bacallao, R., and Wickner, W. (1987). Multiple methods of visualizing the yeast vacuole permit evaluation of its morphology and inheritance during the cell cycle. J. Cell Biol. 105, 1539–47. doi: 10.1083/jcb.105.4.1539.

28. Chung, J.H., Lester, R.L., and Dickson, R.C. (2003). Sphingolipid requirement for generation of a functional v1 component of the vacuolar ATPase. J. Biol. Chem. 278, 28872–81. doi: 10.1074/jbc.M300943200.

29. Odorizzi, G., Babst, M., and Emr, S.D. (1998). Fab1p PtdIns(3)P 5-kinase function essential for protein sorting in the multivesicular body. Cell 95, 847–58. doi: 10.1016/s0092-8674(00)81707-9.

30. Costanzo, M., Baryshnikova, A., Bellay, J., Kim, Y., Spear, E.D., Sevier, C.S., Ding, H., Koh, J.L., Toufighi, K., Mostafavi, S., Prinz, J., St Onge, R.P., VanderSluis, B., Makhnevych, T., Vizeacoumar, F.J., Alizadeh, S., Bahr, S., Brost, R.L., Chen, Y., Cokol, M., Deshpande, R., Li, Z., Lin, Z.Y., Liang, W., Marback, M., Paw, J., San Luis, B. J., Shuteriqi, E., Tong, A.H., van Dyk, N., Wallace, I.M., Whitney, J.A., Weirauch, M.T., Zhong, G., Zhu, H., Houry, W.A., Brudno, M., Ragibizadeh, S., Papp, B., Pál, C., Roth, F.P., Giaever, G., Nislow, C., Troyanskaya, O.G., Bussey, H., Bader, G.D., Gingras, A.C., Morris, Q.D., Kim, P.M., Kaiser, C.A., Myers, C.L., Andrews, B.J., and Boone, C. (2010). The genetic landscape of a cell. Science 327, 425–31. doi: 10.1126/science.1180823.

31. Sharifpoor, S., van Dyk, D., Costanzo, M., Baryshnikova, A., Friesen, H., Douglas, A.C., Youn, J.Y., VanderSluis, B., Myers, C.L., Papp, B., Boone, C., and Andrews, B.J. (2012). Functional wiring of the yeast kinome revealed by global analysis of genetic network motifs. Genome Res. 22, 791–801. doi: 10.1101/gr.129213.111.

32. Voynova, N.S., Roubaty, C., Vazquez, H.M., Mallela, S.K., Ejsing, C.S., and Conzelmann, A. (2015). Saccharomyces cerevisiae Is Dependent on Vesicular Traffic between the Golgi Apparatus and the Vacuole When Inositolphosphorylceramide Synthase Aur1 Is Inactivated. Eukaryot. Cell. 14, 1203–16. doi: 10.1128/EC.00117-15.

33. Suzuki, K., Kirisako, T., Kamada, Y., Mizushima, N., Noda, T., and Ohsumi, Y. (2001). The pre-autophagosomal structure organized by concerted functions of APG genes is essential for autophagosome formation. EMBO J. 20, 5971–81. doi: 10.1093/emboj/20.21.5971.

34. Kamada, Y., Funakoshi, T., Shintani, T., Nagano, K., Ohsumi, M., and Ohsumi, Y. (2000). Tor-mediated induction of autophagy via an Apg1 protein kinase complex. J. Cell Biol. 150, 1507–13. doi: 10.1083/jcb.150.6.1507.

35. Bernard, A., Jin, M., Xu, Z., and Klionsky, D.J. (2015). A large-scale analysis of autophagy-related gene expression identifies new regulators of autophagy. Autophagy 11, 2114–2122. doi: 10.1080/15548627.2015.1099796.

36. Hu, G., McQuiston, T., Bernard, A., Park, Y.D., Qiu, J., Vural, A., Zhang, N., Waterman, S.R., Blewett, N.H., Myers, T.G., Maraia, R.J., Kehrl, J.H., Uzel, G., Klionsky, D.J., and Williamson, P.R. (2015). A conserved mechanism of TOR-dependent RCK-mediated mRNA degradation regulates autophagy. Nat. Cell Biol. 17, 930–942. doi: 10.1038/ncb3189.

37. Huber, A., French, S.L., Tekotte, H., Yerlikaya, S., Stahl, M., Perepelkina, M.P., Tyers, M., Rougemont, J., Beyer, A.L., and Loewith, R. (2011). Sch9 regulates ribosome biogenesis via Stb3, Dot6 and Tod6 and the histone deacetylase complex RPD3L. EMBO J. 30, 3052–64. doi: 10.1038/emboj.2011.221.

38. Lei, Y., Huang, Y., Wen, X., Yin, Z., Zhang, Z., and Klionsky, D.J. (2022). How Cells Deal with the Fluctuating Environment: Autophagy Regulation under Stress in Yeast and Mammalian Systems. Antioxidants (Basel) 11, 304. doi: 10.3390/antiox11020304.

39. Lippman, S.I., and Broach, J.R. (2009). Protein kinase A and TORC1 activate genes for ribosomal biogenesis by inactivating repressors encoded by Dot6 and its homolog Tod6. Proc. Natl. Acad. Sci. USA 106, 19928–33. doi: 10.1073/pnas.0907027106.

40. Reinke, A., Chen, J.C., Aronova, S., and Powers, T. (2006). Caffeine targets TOR complex I and provides evidence for a regulatory link between the FRB and kinase domains of Tor1p. J. Biol. Chem. 281, 31616–26. doi: 10.1074/jbc.M603107200.

41. Shamji, A.F., Kuruvilla, F.G., and Schreiber, S.L. (2000). Partitioning the transcriptional program induced by rapamycin among the effectors of the Tor proteins. Curr. Biol. 10, 1574–81. doi: 10.1016/s0960-9822(00)00866-6.

42. Zurita-Martinez, S.A., and Cardenas, M.E. (2005). Tor and cyclic AMP-protein kinase A: two parallel pathways regulating expression of genes required for cell growth. Eukaryot. Cell 4, 63–71. doi: 10.1128/EC.4.1.63-71.2005.

43. Bonawitz, N.D., Chatenay-Lapointe, M., Pan, Y., and Shadel, G.S. (2007). Reduced TOR signalling extends chronological life span via increased respiration and upregulation of mitochondrial gene expression. Cell Metab. 5, 265–77. doi: 10.1016/j.cmet.2007.02.009.

44. Kaeberlein, M., Powers, R.W. 3rd, Steffen, K.K., Westman, E.A., Hu, D., Dang, N., Kerr, E.O., Kirkland, K.T., Fields, S., and Kennedy, B.K. (2005). Regulation of yeast replicative life span by TOR and Sch9 in response to nutrients. Science 310, 1193–6. doi: 10.1126/science.1115535.

45. Wei, M., Fabrizio, P., Madia, F., Hu, J., Ge, H., Li, L.M., and Longo, V.D. (2009). Tor1/Sch9-regulated carbon source substitution is as effective as calorie restriction in life span extension. PLoS Genet. 5, e1000467. doi: 10.1371/journal.pgen.1000467.

46. Ungar, L., Harari, Y., Toren, A., and Kupiec, M. (2011). Tor complex 1 controls telomere length by affecting the level of Ku. Curr. Biol. 21, 2115–20. doi: 10.1016/j.cub.2011.11.024.

47. Urban, J., Soulard, A., Huber, A., Lippman, S., Mukhopadhyay, D., Deloche, O., Wanke, V., Anrather, D., Ammerer, G., Riezman, H., Broach, J.R., De Virgilio, C., Hall, M.N., and Loewith, R. (2007). Sch9 is a major target of TORC1 in Saccharomyces cerevisiae. Mol. Cell 26, 663–74. doi: 10.1016/j.molcel.2007.04.020.

48. Horvath, A., Sütterlin, C., Manning-Krieg, U., Movva, N.R., and Riezman, H. (1994). Ceramide synthesis enhances transport of GPI-anchored proteins to the Golgi apparatus in yeast. EMBO J. 13, 3687–95. doi: 10.1002/j.1460-2075.1994.tb06678.x

49. Powis, K., Zhang, T., Panchaud, N., Wang, R., De Virgilio, C., and Ding, J. (2015). Crystal structure of the Ego1-Ego2-Ego3 complex and its role in promoting Rag GTPase-dependent TORC1 signalling. Cell Res. 25, 1043–59. doi: 10.1038/cr.2015.86.

50. Kira, S., Kumano, Y., Ukai, H., Takeda, E., Matsuura, A., and Noda, T. (2016). Dynamic relocation of the TORC1-Gtr1/2-Ego1/2/3 complex is regulated by Gtr1 and Gtr2. Mol. Biol. Cell 27, 382–96. doi: 10.1091/mbc.E15-07-0470.

51. Zeng, Q., Araki, Y., and Noda, T. (2024). Pib2 is a cysteine sensor involved in TORC1 activation in Saccharomyces cerevisiae. Cell Rep. 43, 113599. doi: 10.1016/j.celrep.2023.113599.

52. Dickson, R.C. (2010). Roles for sphingolipids in Saccharomyces cerevisiae. Adv. Exp. Med. Biol. 688, 217–31. doi: 10.1007/978-1-4419-6741-1_15.

53. Esch, B.M., Limar, S., Bogdanowski, A., Gournas, C., More, T., Sundag, C., Walter, S., Heinisch, J.J., Ejsing, C.S., André, B., and Fröhlich, F. (2020). Uptake of exogenous serine is important to maintain sphingolipid homeostasis in Saccharomyces cerevisiae. PLoS Genet. 16, e1008745. doi: 10.1371/journal.pgen.1008745.

54. Hepowit, N.L., Macedo, J.K.A., Young, L.E.A., Liu, K., Sun, R.C., MacGurn, J.A., and Dickson, R.C. (2021). Enhancing lifespan of budding yeast by pharmacological lowering of amino acid pools. Aging (Albany NY) 13, 7846–7871. doi: 10.18632/aging.202849.

55. Subramanian, K., Dietrich, L.E., Hou, H., LaGrassa, T.J., Meiringer, C.T., and Ungermann, C. (2006). Palmitoylation determines the function of Vac8 at the yeast vacuole. J. Cell Sci. 119, 2477–85. doi: 10.1242/jcs.02972.

56. Epstein, S., Castillon, G.A., Qin, Y., and Riezman, H. (2012). An essential function of sphingolipids in yeast cell division. Mol. Microbiol. 84, 1018–32. doi: 10.1111/j.1365-2958.2012.08087.x.

57. Rodriguez-Gallardo, S., Kurokawa, K., Sabido-Bozo, S., Cortes-Gomez, A., Ikeda, A., Zoni, V., Aguilera-Romero, A., Perez-Linero, A.M., Lopez, S., Waga, M., Araki, M., Nakano, M., Riezman, H., Funato, K., Vanni, S., Nakano, A., and Muñiz, M. (2020). Ceramide chain length-dependent protein sorting into selective endoplasmic reticulum exit sites. Sci. Adv. 6, eaba8237. doi: 10.1126/sciadv.aba8237.

58. Schlarmann, P., Hanaoka, K., Ikeda, A., Muñiz, M., and Funato, K. (2024). Ceramide sorting into non-vesicular transport is independent of acyl chain length in budding yeast. Biochem. Biophys. Res. Commun. 715, 149980. doi: 10.1016/j.bbrc.2024.149980.

59. Pinto, S.N., Silva, L.C., Futerman, A.H., and Prieto, M. (2011). Effect of ceramide structure on membrane biophysical properties: the role of acyl chain length and unsaturation. Biochim. Biophys. Acta 1808, 2753–60. doi: 10.1016/j.bbamem.2011.07.023.

60. Parasassi, T., and Gratton, E. (1995). Membrane lipid domains and dynamics as detected by Laurdan fluorescence. J. Fluoresc. 5, 59–69. doi: 10.1007/BF00718783.

61. Klose, C., Ejsing, C.S., García-Sáez, A.J., Kaiser, H.J., Sampaio, J.L., Surma, M.A., Shevchenko, A., Schwille, P., and Simons, K. (2010). Yeast lipids can phase-separate into micrometer-scale membrane domains. J. Biol. Chem. 285, 30224–32. doi: 10.1074/jbc.M110.123554.

62. Aronova, S., Wedaman, K., Aronov, P.A., Fontes, K., Ramos, K., Hammock, B.D., and Powers, T. (2008). Regulation of ceramide biosynthesis by TOR complex 2. Cell Metab. 7, 148–58. doi: 10.1016/j.cmet.2007.11.015.

63. Olson, D.K., Fröhlich, F., Farese, R.V. Jr, and Walther, T.C. (2016). Taming the sphinx: Mechanisms of cellular sphingolipid homeostasis. Biochim. Biophys. Acta 1861, 784–792. doi: 10.1016/j.bbalip.2015.12.021.

64. Niles, B.J., Joslin, A.C., Fresques, T., and Powers, T. (2014). TOR complex 2-Ypk1 signalling maintains sphingolipid homeostasis by sensing and regulating ROS accumulation. Cell Rep. 6, 541–52. doi: 10.1016/j.celrep.2013.12.040.

65. Niles, B.J., Mogri, H., Hill, A., Vlahakis, A., and Powers T. (2012). Plasma membrane recruitment and activation of the AGC kinase Ypk1 is mediated by target of rapamycin complex 2 (TORC2) and its effector proteins Slm1 and Slm2. Proc. Natl. Acad. Sci. USA 109, 1536–41. doi: 10.1073/pnas.1117563109.

66. Roelants, F.M., Breslow, D.K., Muir, A., Weissman, J.S., and Thorner, J. (2011). Protein kinase Ypk1 phosphorylates regulatory proteins Orm1 and Orm2 to control sphingolipid homeostasis in Saccharomyces cerevisiae. Proc. Natl. Acad. Sci. USA 108, 19222–7. doi: 10.1073/pnas.1116948108.

67. Muir, A., Ramachandran, S., Roelants, F.M., Timmons, G., and Thorner, J. (2014). TORC2-dependent protein kinase Ypk1 phosphorylates ceramide synthase to stimulate synthesis of complex sphingolipids. Elife 3, e03779. doi: 10.7554/eLife.03779.

68. Gao, J., Nicastro, R., Péli-Gulli, M.P., Grziwa, S., Chen, Z., Kurre, R., Piehler, J., De Virgilio, C., Fröhlich, F., and Ungermann, C. (2022). The HOPS tethering complex is required to maintain signalling endosome identity and TORC1 activity. J. Cell Biol. 221, e202109084. doi: 10.1083/jcb.202109084.

69. Kingsbury, J.M., Sen, N.D., Maeda, T., Heitman, J., and Cardenas, M.E. (2014). Endolysosomal membrane trafficking complexes drive nutrient-dependent TORC1 signalling to control cell growth in Saccharomyces cerevisiae. Genetics 196, 1077–89. doi: 10.1534/genetics.114.161646.

70. Takeda, E., Jin, N., Itakura, E., Kira, S., Kamada, Y., Weisman, L.S., Noda, T., and Matsuura, A. (2018). Vacuole-mediated selective regulation of TORC1-Sch9 signalling following oxidative stress. Mol. Biol. Cell 29, 510–522. doi: 10.1091/mbc.E17-09-0553.

71. Zurita-Martinez, S.A., Puria, R., Pan, X., Boeke, J.D., and Cardenas, M.E. (2007). Efficient Tor signalling requires a functional class C Vps protein complex in Saccharomyces cerevisiae. Genetics 176, 2139–50. doi: 10.1534/genetics.107.072835.

72. Binda, M., Péli-Gulli, M.P., Bonfils, G., Panchaud, N., Urban, J., Sturgill, T.W., Loewith, R., and De Virgilio, C. (2009). The Vam6 GEF controls TORC1 by activating the EGO complex. Mol. Cell 35, 563–73. doi: 10.1016/j.molcel.2009.06.033.

73. Guri, Y., Colombi, M., Dazert, E., Hindupur, S.K., Roszik, J., Moes, S., Jenoe, P., Heim, M.H., Riezman, I., Riezman, H., and Hall, M.N. (2017). mTORC2 Promotes Tumorigenesis via Lipid Synthesis. Cancer Cell 32, 807–823.e12. doi: 10.1016/j.ccell.2017.11.011

74. Zhang, L., Li, Y., Wang, Y., Qiu, Y., Mou, H., Deng, Y., Yao, J., Xia, Z., Zhang, W., Zhu, D., Qiu, Z., Lu, Z., Wang, J., Yang, Z., Mao, G., Chen, D., Sun, L., Liu, L., and Ju, Z. (2022). mTORC2 Facilitates Liver Regeneration Through Sphingolipid-Induced PPAR-α-Fatty Acid Oxidation. Cell. Mol. Gastroenterol. Hepatol. 14, 1311–1331. doi: 10.1016/j.jcmgh.2022.07.011.

75. Zhu, H., Shen, H., Sewell, A.K., Kniazeva, M, and Han, M. (2013). A novel sphingolipid-TORC1 pathway critically promotes postembryonic development in Caenorhabditis elegans. Elife. 2, e00429. doi: 10.7554/eLife.00429.

76. Zhu, M., Teng, F., Li, N., Zhang, L., Zhang, S., Xu, F., Shao, J., Sun, H., and Zhu, H. (2021). Monomethyl branched-chain fatty acid mediates amino acid sensing upstream of mTORC1. Dev. Cell 56, 2692–2702.e5. doi: 10.1016/j.devcel.2021.09.010.

77. Kinghorn, K.J., Grönke, S., Castillo-Quan, J.I., Woodling, N.S., Li, L., Sirka, E., Gegg, M., Mills, K., Hardy, J., Bjedov, I., and Partridge, L. (2016). A Drosophila Model of Neuronopathic Gaucher Disease Demonstrates Lysosomal-Autophagic Defects and Altered mTOR Signalling and Is Functionally Rescued by Rapamycin. J. Neurosci. 36, 11654–11670. doi: 10.1523/JNEUROSCI.4527-15.2016.

78. Magalhaes, J., Gegg, M.E., Migdalska-Richards, A., Doherty, M.K., Whitfield, P.D., and Schapira, A.H. (2016). Autophagic lysosome reformation dysfunction in glucocerebrosidase deficient cells: relevance to Parkinson disease. Hum. Mol. Genet. 25, 3432–3445. doi: 10.1093/hmg/ddw185.

79. Wang, F., Dai, Y., Zhu, X., Chen, Q., Zhu, H., Zhou, B., Tang, H., and Pang, S. (2021). Saturated very long chain fatty acid configures glycosphingolipid for lysosome homeostasis in long-lived C. elegans. Nat. Commun. 12, 5073. doi: 10.1038/s41467-021-25398-6.

80. Hartwig, P., and Höglinger, D. (2021). The Glucosylceramide Synthase Inhibitor PDMP Causes Lysosomal Lipid Accumulation and mTOR Inactivation. Int. J. Mol. Sci. 22, 7065. doi: 10.3390/ijms22137065.

81. Ode, T., Podyma-Inoue, K.A., Terasawa, K., Inokuchi, J.I., Kobayashi, T., Watabe, T., Izumi, Y., and Hara-Yokoyama, M. (2017). PDMP, a ceramide analogue, acts as an inhibitor of mTORC1 by inducing its translocation from lysosome to endoplasmic reticulum. Exp. Cell Res. 350, 103–114. doi: 10.1016/j.yexcr.2016.11.011.

82. Castellano, B.M., Thelen, A.M., Moldavski, O., Feltes, M., van der Welle, R.E., Mydock-McGrane, L., Jiang, X., van Eijkeren, R.J., Davis, O.B., Louie, S.M., Perera, R.M., Covey, D.F., Nomura, D.K., Ory, D.S., and Zoncu, R. (2017). Lysosomal cholesterol activates mTORC1 via an SLC38A9-Niemann-Pick C1 signalling complex. Science 355, 1306–1311. doi: 10.1126/science.aag1417.

83. Xu, J., Dang, Y., Ren, Y.R., and Liu, J.O. (2010). Cholesterol trafficking is required for mTOR activation in endothelial cells. Proc. Natl. Acad. Sci. USA 107, 4764–9. doi: 10.1073/pnas.0910872107.

84. Nada, S., Hondo, A., Kasai, A., Koike, M., Saito, K., Uchiyama, Y., and Okada, M. (2009). The novel lipid raft adaptor p18 controls endosome dynamics by anchoring the MEK-ERK pathway to late endosomes. EMBO J. 28, 477–89. doi: 10.1038/emboj.2008.308.

85. Fang, Y., Vilella-Bach, M., Bachmann, R., Flanigan, A., and Chen, J. (2001). Phosphatidic acid-mediated mitogenic activation of mTOR signalling. Science 294, 1942–5. doi: 10.1126/science.1066015.

86. Veverka, V., Crabbe, T., Bird, I., Lennie, G., Muskett, F.W., Taylor, R.J., and Carr, M.D. (2008). Structural characterization of the interaction of mTOR with phosphatidic acid and a novel class of inhibitor: compelling evidence for a central role of the FRB domain in small molecule-mediated regulation of mTOR. Oncogene 27, 585–95. doi: 10.1038/sj.onc.1210693.

87. Frias, M.A., Hatipoglu, A., and Foster, D.A. (2023). Regulation of mTOR by phosphatidic acid. Trends Endocrinol. Metab. 34, 170–180. doi: 10.1016/j.tem.2023.01.004.

88. Frias, M.A., Mukhopadhyay, S., Lehman, E., Walasek, A., Utter, M., Menon, D., and Foster, D.A. (2020). Phosphatidic acid drives mTORC1 lysosomal translocation in the absence of amino acids. J. Biol. Chem. 295, 263–274. doi: 10.1074/jbc.RA119.010892.

89. Guo, Z., Zhao, K., Feng, X., Yan, D., Yao, R., Chen, Y., Bao, L., and Wang, Z. (2019). mTORC2 Regulates Lipogenic Gene Expression through PPARγ to Control Lipid Synthesis in Bovine Mammary Epithelial Cells. Biomed Res. Int. 2019, 5196028. doi: 10.1155/2019/5196028.

90. Battaglioni, S., Benjamin, D., Wälchli, M., Maier, T., and Hall, M.N. (2022). mTOR substrate phosphorylation in growth control. Cell 185, 1814–1836. doi: 10.1016/j.cell.2022.04.013.

91. Le Bacquer, O., Queniat, G., Gmyr, V., Kerr-Conte, J., Lefebvre, B., and Pattou, F. (2013). mTORC1 and mTORC2 regulate insulin secretion through Akt in INS-1 cells. J. Endocrinol. 216, 21–9. doi: 10.1530/JOE-12-0351.

92. Kato, M., and Wickner, W. (2001). Ergosterol is required for the Sec18/ATP-dependent priming step of homotypic vacuole fusion. EMBO J. 20, 4035–40. doi: 10.1093/emboj/20.15.4035.

93. Lang, T., Bruns, D., Wenzel, D., Riedel, D., Holroyd, P., Thiele, C., and Jahn, R. (2001). SNAREs are concentrated in cholesterol-dependent clusters that define docking and fusion sites for exocytosis. EMBO J. 20, 2202–13. doi: 10.1093/emboj/20.9.2202.

94. Heese-Peck, A., Pichler, H., Zanolari, B., Watanabe, R., Daum, G., and Riezman, H. (2002). Multiple functions of sterols in yeast endocytosis. Mol. Biol. Cell 13, 2664–80. doi: 10.1091/mbc.e02-04-0186.

95. Zhang, Y.Q., Gamarra, S., Garcia-Effron, G., Park, S., Perlin, D.S., and Rao, R. (2010). Requirement for ergosterol in V-ATPase function underlies antifungal activity of azole drugs. PLoS Pathog. 6, e1000939. doi: 10.1371/journal.ppat.1000939.

96. Tani, M., and Toume, M. (2015). Alteration of complex sphingolipid composition and its physiological significance in yeast Saccharomyces cerevisiae lacking vacuolar ATPase. Microbiology (Reading) 161, 2369–83. doi: 10.1099/mic.0.000187.

97. Coonrod, E.M., Graham, L.A., Carpp, L.N., Carr, T.M., Stirrat, L., Bowers, K., Bryant, N.J., and Stevens TH. (2013). Homotypic vacuole fusion in yeast requires organelle acidification and not the V-ATPase membrane domain. Dev. Cell 27, 462–8. doi: 10.1016/j.devcel.2013.10.014.

98. Desfougères, Y., Vavassori, S., Rompf, M., Gerasimaite, R., and Mayer, A. (2016). Organelle acidification negatively regulates vacuole membrane fusion in vivo. Sci. Rep. 6, 29045. doi: 10.1038/srep29045.

99. Brett, C.L., Kallay, L., Hua, Z., Green, R., Chyou, A., Zhang, Y., Graham, T.R., Donowitz, M., and Rao, R. (2011). Genome-wide analysis reveals the vacuolar pH-stat of Saccharomyces cerevisiae. PLoS One 6, e17619. doi: 10.1371/journal.pone.0017619.

100. Liu, Y., Wang, R., Liu, J., Fan, M., Ye, Z., Hao, Y., Xie, F., Wang, T., Jiang, Y., Liu, N., Cui, X., Lv, Q., and Yan, L. (2024). The vacuolar fusion regulated by HOPS complex promotes hyphal initiation and penetration in Candida albicans. Nat. Commun. 15, 4131. doi: 10.1038/s41467-024-48525-5.

101. Alfatah, M., Cui, L., Goh, C.J.H., Cheng, T.Y.N., Zhang, Y., Naaz, A., Wong, J.H., Lewis, J., Poh, W.J., and Arumugam, P. (2023). Metabolism of glucose activates TORC1 through multiple mechanisms in Saccharomyces cerevisiae. Cell Rep. 42, 113205. doi: 10.1016/j.celrep.2023.113205.

102. Jansen, G., Wu, C., Schade, B., Thomas, D.Y., and Whiteway, M. (2005). Drag&Drop cloning in yeast. Gene 344, 43–51. doi: 10.1016/j.gene.2004.10.016.

103. Sherman, F., Fink, G.R., and Lawrence, C. (1979). Methods in Yeast Genetics. Cold Spring Harbor Laboratory Press, Cold Spring Harbor, New York.

104. Smith, D.L., Jr, McClure, J.M., Matecic, M., and Smith, J.S. (2007). Calorie restriction extends the chronological lifespan of Saccharomyces cerevisiae independently of the Sirtuins. Aging Cell 6, 649–62. doi: 10.1111/j.1474-9726.2007.00326.x.

105. LaGrassa, T.J., and Ungermann, C. (2005). The vacuolar kinase Yck3 maintains organelle fragmentation by regulating the HOPS tethering complex. J. Cell Biol. 168, 401–14. doi: 10.1083/jcb.200407141.

106. Wu, H., Zheng, X., Araki, Y., Sahara, H., Takagi, H., and Shimoi, H. (2006). Global gene expression analysis of yeast cells during sake brewing. Appl. Environ. Microbiol. 72, 7353–8. doi: 10.1128/AEM.01097-06.

107. Craven, R.J., and Petes, T.D. (1999). Dependence of the regulation of telomere length on the type of subtelomeric repeat in the yeast Saccharomyces cerevisiae. Genetics 152, 1531–41. doi: 10.1093/genetics/152.4.1531.

108. Yabuki, Y., Ikeda, A., Araki, M., Kajiwara, K., Mizuta, K., and Funato K. (2019). Sphingolipid/Pkh1/2-TORC1/Sch9 Signalling Regulates Ribosome Biogenesis in Tunicamycin-Induced Stress Response in Yeast. Genetics 212, 175–186. doi: 10.1534/genetics.118.301874.

109. Ikeda, A., Hanaoka, K., and Funato K. (2021). Protocol for measuring sphingolipid metabolism in budding yeast. STAR Protoc. 2, 100412. doi: 10.1016/j.xpro.2021.100412.

110. Kajiwara, K., Ikeda, A., Aguilera-Romero, A., Castillon, G.A., Kagiwada, S., Hanada, K., Riezman, H., Muñiz, M., and Funato, K. (2014). Osh proteins regulate COPII-mediated vesicular transport of ceramide from the endoplasmic reticulum in budding yeast. J. Cell Sci. 127, 376–87. doi: 10.1242/jcs.132001.

111. Chahomchuen, T., Hondo, K., Ohsaki, M., Sekito, T., and Kakinuma, Y. (2009). Evidence for Avt6 as a vacuolar exporter of acidic amino acids in Saccharomyces cerevisiae cells. J. Gen. Appl. Microbiol. 55, 409–17. doi: 10.2323/jgam.55.409.

112. Adhyapak, P., Srivatsav, A.T., Mishra, M., Singh, A., Narayan, R., and Kapoor, S. (2020). Dynamical Organization of Compositionally Distinct Inner and Outer Membrane Lipids of Mycobacteria. Biophys. J. 118, 1279–1291. doi: 10.1016/j.bpj.2020.01.027.

113. Horigome, C., Okada, T., Matsuki, K., and Mizuta, K. (2008). A ribosome assembly factor Ebp2p, the yeast homolog of EBNA1-binding protein 2, is involved in the secretory response. Biosci. Biotechnol. Biochem. 72, 1080–6. doi: 10.1271/bbb.70817.

114. Forsburg, S.L. (2001). The art and design of genetic screens: yeast. Nat. Rev. Genet. 2, 659–68. doi: 10.1038/35088500

115. Hashida-Okado, T., Ogawa, A., Endo, M., Yasumoto, R., Takesako, K., and Kato, I. (1996). AUR1, a novel gene conferring aureobasidin resistance on Saccharomyces cerevisiae: a study of defective morphologies in Aur1p-depleted cells. Mol. Gen. Genet. 251, 236–44. doi: 10.1007/BF02172923.

