## Supplementary material for "Sphingolipids integrate TORC2 with TORC1 through vacuolar liquid-ordered domain formation": Table S1

**Table S1. Yeast strains used in this study**

| Strain (names or number) | | | Genotype | Source or Reference | Figure |
|---|---|---|---|---|---|
| Wild type | BY4742 | MATα | his3Δ1 leu2Δ0 ura3Δ0 lys2Δ0 | Open biosystem, Inc | 1A, 3I, 4B, 4C, 7A-F, S1, S2A-C, S5B, S12A, S12B |
| Knock out haploid collection | | MATα | his3Δ1 leu2Δ0 ura3Δ0 lys2Δ0 xxx::KanMX | Open biosystem, Inc | 1A, 3I, 4B, 7A-F, S1, S2A-C, S5B, S6A-C, S12A, S12B |
| Wild type | BY4741 | MATa | his3Δ1 leu2Δ0 ura3Δ0 met15Δ0 | Open biosystem, Inc | |
| Wild type | FKY1131 | MATa | his3Δ1 leu2Δ0 ura3Δ0 lys2Δ ; same as BY4741 but MET15 lys2Δ | Kajiwara et al., Mol. Microbiol., 2012 | 1B, 2A, 2B |
| AUR1 | FKY2040 | MATα | his3Δ1 leu2Δ0 ura3Δ0 aur1Δ::HIS3 carrying pRS415/AUR1(LEU2, CEN) | This study | 1C, 1E, 1F, 1I, S4A-D, S5A, S6A-D, S7A |
| aur1-1 | FKY2041 | MATα | his3Δ1 leu2Δ0 ura3Δ0 aur1Δ::HIS3 carrying pRS415/aur1-1(LEU2, CEN) | This study | 1C-F, 1I, S4A-D, S5A, S6A-D, S7A |
| Wild type | RH1800 | MATa | his4 leu2 ura3 bar1 | Zanolari et al., EMBO J., 2000 | 1G, 3H, S6A, S6C |
| lcb1-100 | RH3802 | MATa | his3 his4 leu2 ura3 lys2 ade2 bar1 lcb1-100 | Friant et al., EMBO J., 2000 | 1D, 1G, 3H, S6A-C |
| pep4Δ AUR1 | FKY2568 | MATα | his3Δ1 leu2Δ0 ura3Δ0 pep4Δ::KanMX aur1Δ::HIS3 carrying pRS415/AUR1(LEU2, CEN) | This study | 1H |
| pep4Δ aur1-1 | FKY2569 | MATα | his3Δ1 leu2Δ0 ura3Δ0 pep4Δ::KanMX aur1Δ::HIS3 carrying pRS415/aur1-1(LEU2, CEN) | This study | 1H |
| AUR1 | FKY1189-1 | MATa | his3Δ1 leu2Δ0 ura3Δ0 lys2Δ0 met15Δ0 aur1Δ::HIS3 carrying pRS415/AUR1(LEU2, CEN) | This study | 2C, 3A, 3C, 3D, 4A, 4B, 4D, 4E |
| aur1-1 | FKY1189-2 | MATa | his3Δ1 leu2Δ0 ura3Δ0 lys2Δ0 met15Δ0 aur1Δ::HIS3 carrying pRS415/aur1-1(LEU2, CEN) | This study | 2C, 3A, 3C, 4A, 4B, 4D, 4E |
| aur1-6 | FKY1189-3 | MATa | his3Δ1 leu2Δ0 ura3Δ0 lys2Δ0 met15Δ0 aur1Δ::HIS3 carrying pRS415/aur1-6(LEU2, CEN) | This study | 2C, 3A |
| aur1-18 | FKY1189-4 | MATa | his3Δ1 leu2Δ0 ura3Δ0 lys2Δ0 met15Δ0 aur1Δ::HIS3 carrying pRS415/aur1-18(LEU2, CEN) | This study | 2C, 3A, 3C, 4A |
| AUR1 | FKY2524 | MATa | his3-11,15 ade2-1 ura3-1 leu2-3,112 trp1-1 can1-100 aur1Δ::HIS3 carrying pRS415/AUR1(LEU2, CEN) | This study | 2D-F, 3A, 3B, S7B, S10F |
| aur1-1 | FKY2526 | MATa | his3-11,15 ade2-1 ura3-1 leu2-3,112 trp1-1 can1-100 aur1Δ::HIS3 carrying pRS415/aur1-1(LEU2, CEN) | This study | 2D-F, 3A, 3B, S7B, S10F |
| aur1-6 | FKY2528 | MATa | his3-11,15 ade2-1 ura3-1 leu2-3,112 trp1-1 can1-100 aur1Δ::HIS3 carrying pRS415/aur1-6(LEU2, CEN) | This study | 3A, 3B, S7B |
| aur1-18 | FKY2530 | MATa | his3-11,15 ade2-1 ura3-1 leu2-3,112 trp1-1 can1-100 aur1Δ::HIS3 carrying pRS415/aur1-18(LEU2, CEN) | This study | 3A, 3B, S7B |
| AUR1 tor1Δ | FKY2532 | MATa | his3-11,15 ade2-1 ura3-1 leu2-3,112 trp1-1 can1-100 tor1Δ::TRP1 aur1Δ::HIS3 carrying pRS415/AUR1(LEU2, CEN) | This study | 2E, 3B, S7B |
| aur1-1 tor1Δ | FKY2534 | MATa | his3-11,15 ade2-1 ura3-1 leu2-3,112 trp1-1 can1-100 tor1Δ::TRP1 aur1Δ::HIS3 carrying pRS415/aur1-1(LEU2, CEN) | This study | 3B, S7B |
| aur1-6 tor1Δ | FKY2536 | MATa | his3-11,15 ade2-1 ura3-1 leu2-3,112 trp1-1 can1-100 tor1Δ::TRP1 aur1Δ::HIS3 carrying pRS415/aur1-6(LEU2, CEN) | This study | 3B, S7B |
| aur1-18 tor1Δ | FKY2538 | MATa | his3-11,15 ade2-1 ura3-1 leu2-3,112 trp1-1 can1-100 tor1Δ::TRP1 aur1Δ::HIS3 carrying pRS415/aur1-18(LEU2, CEN) | This study | 3B, S7B |
| AUR1 tco89Δ | FKY2611 | MATa | his3-11,15 ade2-1 ura3-1 leu2-3,112 trp1-1 can1-100 tco89Δ::TRP1 aur1Δ::HIS3 carrying pRS415/AUR1(LEU2, CEN) | This study | 3B, S7B |
| aur1-1 tco89Δ | FKY2612 | MATa | his3-11,15 ade2-1 ura3-1 leu2-3,112 trp1-1 can1-100 tco89Δ::TRP1 aur1Δ::HIS3 carrying pRS415/aur1-1(LEU2, CEN) | This study | 3B, S7B |
| Wild type | FKY2603 | MATα | his3 his4 leu2 ura3 lys2 ade2 trp1 bar1 | This study | 3E-G, S10F, S10I |
| lcb1-100 | FKY2604 | MATα | his3 his4 leu2 ura3 lys2 ade2 bar1 lcb1-100 | This study | 3E-G, S10F, S10I |
| tor1Δ | FKY2605 | MATα | his3 his4 leu2 ura3 lys2 ade2 trp1 bar1 tor1Δ::LEU2 | This study | 3E-G |
| lcb1-100 tor1Δ | FKY2606 | MATα | his3 his4 leu2 ura3 lys2 ade2 bar1 lcb1-100 tor1Δ::LEU2 | This study | 3E-G |
| csg1Δ csg2Δ | FKY4407 | MATα | his3Δ1, leu2Δ0, lys2Δ0, ura3Δ0, csg1Δ::KanMX, csg2Δ::KanMX | This study | 3I |
| AUR1 GTR1-GBP | FKY4006-1 | MATα | his3Δ1 leu2Δ0 ura3Δ0 lys2Δ0 met15Δ0 GTR1-GBP::HphMX aur1Δ::HIS3 carrying pRS415/AUR1(LEU2, CEN) | This study | 4C |
| aur1-1 GTR1-GBP | FKY4006-2 | MATα | his3Δ1 leu2Δ0 ura3Δ0 lys2Δ0 met15Δ0 GTR1-GBP::HphMX aur1Δ::HIS3 carrying pRS415/aur1-1(LEU2, CEN) | This study | 4C, 4E |
| ego3Δ GTP1-GBP | FKY4442 | MATa | his3Δ1 leu2Δ0 lys2Δ0 ura3Δ0 GTR1-GBP-6HIS-mcherr::HphMX ego3Δ::KanMX | This study | 4C |
| Wild type GTR1-GBP | FKY3872 | MATα | his3Δ1 leu2Δ0 lys2Δ0 ura3Δ0 GTR1-GBP-6HIS-mcherry::HphMX | This study | 4E |
| aur1-1 VAC8-GFP | FKY4010-2 | MATα | his3Δ1 leu2Δ0 ura3Δ0 lys2Δ0 met15Δ0 VAC8-GFP::KanMX aur1Δ::HIS3 carrying pRS415/aur1-1(LEU2, CEN) | This study | 4E |
| aur1-1 GTR1-GBP VAC8-GFP | FKY4009-2 | MATα | his3Δ1 leu2Δ0 ura3Δ0 lys2Δ0 met15Δ0 GTR1-GBP::HphMX VAC8-GFP::KanMX aur1Δ::HIS3 carrying pRS415/aur1-1(LEU2, CEN | This study | 4E |
| aur1-1 GTR1-GBP VPH1-GFP | FKY3895-2 | MATα | his3Δ1 leu2Δ0 ura3Δ0 lys2Δ0 GTR1-GBP::HphMX VPH1-GFP::KanMX aur1Δ::HIS3 carrying pRS415/aur1-1(LEU2, CEN) | This study | 4E |
| Wild type | RH6095 | MATα | leu2 ura3 lys2 | Sasaki et al., FEBS Lett. 2024 | 5A-G |
| Ghlag1 | RH6979 | MATα | lag1Δ::HIS3 lac1Δ::ADE2 TDH3::GhLAG1::TRP1 leu2 ura3 lys2 | Sasaki et al., FEBS Lett. 2024 | 5A-G |
| AVO3 | PLY567 | MATa | his3 leu2 ura3 lys2 trp1 bar1 AVO3-MYC:TRP1 | Aronova et al., Cell Metab., 2008 | 6A-I, S10G, S10H, S11A-C |
| avo3-30 | PLY694 | MATa | his3 leu2 ura3 lys2 trp1 bar1 avo3-30-MYC:TRP1 | Aronova et al., Cell Metab., 2008 | 6A-I, S10G, S10H, S11A-C |
| AVO3 | FKY6783 | MATa | his3 leu2 ura3 ade2 trp1 bar1 AVO3-MYC:TRP1 | This study | 6J-L |
| AVO3 orm1Δ orm2Δ | FKY6784 | MATα | his3 leu2 ura3 ade2 trp1 bar1 AVO3-MYC:TRP1 orm1Δ:HIS3, orm2Δ:LEU2 | This study | 6L |
| avo3-30 | FKY6785 | MATa | his3 leu2 ura3 ade2 trp1 bar1 avo3-30-MYC:TRP1 | This study | 6J-L |
| avo3-30 orm1Δ orm2Δ | FKY6786 | MATα | his3 leu2 ura3 ade2 trp1 bar1 avo3-30-MYC:TRP1 orm1Δ:HIS3, orm2Δ:LEU2 | This study | 6J-L |
| aur1Δ (pGAL/AUR1-HA) | FKY1189 | MATa | his3Δ1 leu2Δ0 ura3Δ0 lys2Δ0 met15Δ0 aur1Δ::HIS3 carrying pGAL/AUR1-3HA (URA3, CEN) | This study | S3A |
| aur1Δ vps3Δ (pGAL/AUR1-HA | FKY2665 | MATa | his3Δ1 leu2Δ0 ura3Δ0 lys2Δ0 met15Δ0 vps3Δ::KanMX aur1Δ::HIS3 carrying pGAL/AUR1-3HA (URA3, CEN) | This study | S3A |
| aur1Δ vma4Δ (pGAL/AUR1-H | FKY2654 | MATa | his3Δ1 leu2Δ0 ura3Δ0 lys2Δ0 met15Δ0 vma4Δ::KanMX aur1Δ::HIS3 carrying pGAL/AUR1-3HA (URA3, CEN) | This study | S3B |
| aur1Δ stp22Δ (pGAL/AUR1-H | FKY2625 | MATa | his3Δ1 leu2Δ0 ura3Δ0 lys2Δ0 stp22Δ::KanMX aur1Δ::HIS3 carrying pGAL/AUR1-3HA (URA3, CEN) | This study | S3C |
| aur1Δ vps36Δ (pGAL/AUR1-H | FKY2627 | MATa | his3Δ1 leu2Δ0 ura3Δ0 lys2Δ0 met15Δ0 vps36Δ::KanMX aur1Δ::HIS3 carrying pGAL/AUR1-3HA (URA3, CEN) | This study | S3C |
| aur1Δ vps20Δ (pGAL/AUR1-H | FKY2634 | MATa | his3Δ1 leu2Δ0 ura3Δ0 lys2Δ0 met15Δ0 vps20Δ::KanMX aur1Δ::HIS3 carrying pGAL/AUR1-3HA (URA3, CEN) | This study | S3C |
| aur1Δ snf7Δ (pGAL/AUR1-HA | FKY2621 | MATa | his3Δ1 leu2Δ0 ura3Δ0 lys2Δ0 met15Δ0 snf7Δ::KanMX aur1Δ::HIS3 carrying pGAL/AUR1-3HA (URA3, CEN) | This study | S3C |
| aur1-6 | FKY2042 | MATα | his3Δ1 leu2Δ0 ura3Δ0 aur1Δ::HIS3 carrying pRS415/aur1-6(LEU2, CEN) | This study | S4A-D, S5A, S7A |
| aur1-18 | FKY2043 | MATα | his3Δ1 leu2Δ0 ura3Δ0 aur1Δ::HIS3 carrying pRS415/aur1-18(LEU2, CEN) | This study | S4A-D, S5A, S7A |
| Wild type | FKY17 | MATa | his4 leu2 ura3 lys2 ade2 bar1; same as RH1800 but lys2 and ade2 | This study | S5B |
| erg6Δ | RH5684 | MATa | his4 leu2 ura3 lys2 bar1erg6Δ::LEU2 | Guan et al., Mol. Biol. Cell, 2009 | S5B |
| AUR1 EGO1-GBP | FKY5443-1 | MATα | his3Δ1 leu2Δ0 ura3Δ0 lys2Δ0 met15Δ0 EGO1-GBP::KanMX aur1Δ::HIS3 carrying pRS415/AUR1(LEU2, CEN) | This study | S8A, S8B |
| aur1-1 EGO1-GBP | FKY5443-2 | MATα | his3Δ1 leu2Δ0 ura3Δ0 lys2Δ0 met15Δ0 EGO1-GBP::KanMX aur1Δ::HIS3 carrying pRS415/aur1-1(LEU2, CEN) | This study | S8A, S8B |
| AUR1 | FKY4426 | MATa | his3Δ1 leu2Δ0 ura3Δ0 lys2Δ0 aur1Δ::HIS3 carrying pRS415/AUR1(LEU2, CEN) | This study | S9A-C |
| aur1-1 | FKY4427 | MATa | his3Δ1 leu2Δ0 ura3Δ0 lys2Δ0 aur1Δ::HIS3 carrying pRS415/ aur1-1(LEU2, CEN) | This study | S9A-C |
| aur1-18 | FKY4429 | MATa | his3Δ1 leu2Δ0 ura3Δ0 lys2Δ0 aur1Δ::HIS3 carrying pRS415/ aur1-18(LEU2, CEN) | This study | S9A-C |
| AUR1 can1Δ | FKY4430 | MATa | his3Δ1 leu2Δ0 ura3Δ0 lys2Δ0 can1Δ::KanMx aur1Δ::HIS3 carrying pRS415/AUR1(LEU2, CEN) | This study | S9A-C |
| aur1-1 can1Δ | FKY4431 | MATa | his3Δ1 leu2Δ0 ura3Δ0 lys2Δ0 can1Δ::KanMx aur1Δ::HIS3 carrying pRS415/ aur1-1(LEU2, CEN) | This study | S9A-C |
| aur1-18 can1Δ | FKY4433 | MATa | his3Δ1 leu2Δ0 ura3Δ0 lys2Δ0 can1Δ::KanMx aur1Δ::HIS3 carrying pRS415/ aur1-18(LEU2, CEN) | This study | S9A-C |
| Wild type | SEY6210 | MATα | ura3-52 his3@200 lys2-801 leu2-3,112 trp1@901 suc2@9 | Tabuchi et al., Mol. Cell Biol., 2006 | S10B-F |
| stt4-4 | AAY102 | MATα | SEY6210 stt4Δ::HIS3 carrying pRS415/stt4-4 (LEU2, CEN) | Tabuchi et al., Mol. Cell Biol., 2006 | S10B |
| mss4-102 | AAY202 | MATα | SEY6210 mss4Δ::HIS3MX6 carrying Ycplac/mss4-102 (LEU2, CEN) | Tabuchi et al., Mol. Cell Biol., 2006 | S10B |
| pkc1-2 | AAY603 | MATα | SEY6210 pkc1Δ::LEU2 carrying Ycp50/pkc1-2 (URA3, CEN) | Tabuchi et al., Mol. Cell Biol., 2006 | S10B |
| rho1 | AAY608 | MATα | SEY6210 rho1Δ::URA3 carrying pRS316/rho1 (URA3, CEN) | Tabuchi et al., Mol. Cell Biol., 2006 | S10B |
| slm1-1 slm2Δ | AAY1622 | MATα | SEY6210 slm1Δ::HIS3 slm2Δ::HIS3 carrying pRS415/slm1-1(LEU2, CEN) | Tabuchi et al., Mol. Cell Biol., 2006 | S10B-D, S10F |
| pkh1 pkh2Δ | MTY29 | MATα | SEY6210 pkh1Δ::HIS3MX6 pkh2Δ::HIS3MX6 carrying pRS415/pkh1(LEU2, CEN) | Omnus et at., Mol. Bio. Cell, 2016 | S10B-D |
| ypk1 ypk2Δ | MTY77 | MATα | SEY6210 ypk1ts::HIS3 ypk2Δ::HIS3MX6 | Omnus et at., Mol. Bio. Cell, 2016 | S10B-D, S10E |
| AVO3 | FKY5951 | MATa | his3 leu2 ura3 lys2 trp1 bar1 AVO3-MYC:TRP1 cnb1Δ::kanMX | This study | S11A-C |
| avo3-30 | FKY5952 | MATa | his3 leu2 ura3 lys2 trp1 bar1 avo3-30-MYC:TRP1 cnb1Δ::kanMX | This study | S11A-C |
| Wild type | FKY5663 | MATα | his3 leu2 ura3 lys2 trp1 bar1 | Schlarmann et al., J. Biol. Chem., 2024 | S13A, S13B |
